# *Helicobacter pylori* allelic variation in cell surface genes influences human exoproteome binding and stomach tissue adherence

**DOI:** 10.64898/2026.03.06.710112

**Authors:** Jacob P. Frick, Nicole D. Sonnert, Jazmine A. Snow, Moira Overly, Hyojik Yang, Megan N. Stoppler, Aaron M. Ring, Robert K. Ernst, Noah W. Palm, Nina R. Salama

**Author notes:** Correspondence: Nina R. Salama, Fred Hutchinson Cancer Center, 1100 Fairview Ave North, Mail Stop C3-168, Seattle, WA 98109.

## Abstract

*Helicobacter pylori,* the primary etiological agent of gastric cancer, requires chronic infection to promote severe disease. Throughout colonization, the bacteria accumulate genetic variation, which can reshape the interaction between host and pathogen. In this study, we probed adherence as one important characteristic of this relationship. We leveraged a panel of *H. pylori* strains, representing both inter- and intra-host diversity, and BASEHIT, a comprehensive barcoded yeast display library of the human exoproteome, to evaluate the human exoprotein binding characteristics of *H. pylori*. We identified a set of lineage-correlated binding phenotypes and a set of polymorphic cell surface-associated loci that we predict to govern heterogeneous binding. We also identified a general increase in gastric tissue adherence during mouse passage and structural modifications in the O-antigen component lipopolysaccharide. Subsequent sequence analyses identified C-terminal repeat length reduction in either *futB* or glycosyltransferase family 25 galactose transferases as sufficient to alter lipopolysaccharide. Shorter repeat variants were favored during both acute and chronic colonization and conferred a colonization advantage in coinfection. Our data indicate that shorter enzymes, and the resulting shorter O-antigen repeats, are favored during infection. We hypothesize that repeat length polymorphisms modulate enzyme efficiency and disrupt the balance of competing reactions required for O-antigen synthesis.

**Importance:** In this study, we used a high throughput assay to evaluate the adherence of genetically diverse *Helicobacter pylori* isolates to human proteins they may encounter in the stomach. We showed that more closely related *H. pylori* strains bind human proteins similarly. We linked changes in bacterial binding to genetic differences affecting cell surface components. In particular, *H. pylori* strains passaged through mice exhibited increased adherence to stomach tissue, which we associated with structural variation in the enzymes responsible for synthesizing the O-antigen portion of lipopolysaccharide. We also showed that shorter enzyme variants produce distinct O-antigen structures that enhance colonization of the mouse stomach.

## Introduction

*Helicobacter pylori* is a robust colonizer of the human stomach, present in ∼40% of the global population (**1**). While most of those infected will remain asymptomatic with only mild gastritis, a subset will exhibit more severe pathologies, such as peptic ulcer disease, atrophy, and gastric cancer. Infection with *H. pylori* is the main risk factor for developing gastric cancer, which requires a chronic interaction between the bacterium and host to occur (**2, 3**). Our understanding of the factors necessary for *H. pylori* to colonize and cause disease is complicated by the high degree of genetic diversity amongst strains from different sources and even those within a single individual (**4–10**). In the current study, we focus on *H. pylori* cell surface-associated genes as modulators of interactions with host tissue. These genes are enriched for variation during chronic infection and can directly affect the interaction of host and pathogen (**8, 9, 11–14**).

*H. pylori* encodes approximately 60 outer membrane proteins (OMPs), with individual strains expressing unique combinations that define their surface composition (**4, 11, 15**). Many *H. pylori* OMPs are adhesins or have homology to adhesins, which promote adherence to gastric tissue by binding to glycan or peptide epitopes on host cells, mucins, and other bacteria (**11**). The best characterized adhesins are BabA, which binds to blood-group antigens (**16, 17**), SabA, which binds to sialyl-Lewis antigens (**18**), and HopQ, which binds CEACAMs (**19**); however, many remain uncharacterized. During chronic infection, adhesins are known to accumulate genetic variation, which likely affects their functions (**8, 20–22**).

Adhesins are not the only factor contributing to *H. pylori* interaction with host tissue and mucins. The lipopolysaccharide (LPS) of *H. pylori* influences adherence, primarily through display of epitopes (such as Lewis antigens) bound by host lectins (**23–26**), although the O-antigen of LPS has also been implicated as an adhesin itself (**25, 27–29**). A recent body of literature better elucidated the LPS structure of *H. pylori* strain G27 and defined many of the glycosyltransferases involved in its synthesis (**30–32**). Importantly, many of these genes have the capacity for phase variation, which may diversify their expression within an infecting bacterial population (**33–35**). Some LPS glycosyltransferases, namely the FucT fucosyltransferases, also include simple repeats in their C-termini, which allow for elongation or shortening of the protein product, believed to affect its enzymatic activity and the resulting LPS structure (**33, 36–38**). However, the functional impacts of LPS and LPS-gene diversity on *H. pylori-*gastric tissue interactions are not as well characterized.

In this study, we employed an unbiased, comprehensive approach to investigate how *H. pylori* diversity influences host interactions. We leveraged the J99 culture collection of *H. pylori* isolates obtained from six years of chronic infection of a single host (**8, 10**). These isolates, representing four distinct phylogenetic subgroups and two disease states, along with several commonly used reference strains, were analyzed using a combination of high-throughput human exoprotein adhesion profiling, murine stomach colonization experiments, and *ex vivo* adherence assays to assess functional variation in *H. pylori* cell surface interactions. This analysis identified 92 human proteins with *H. pylori* interactions and a set of polymorphic bacterial genetic loci associated with binding phenotypes. We showed that mouse passage of human *H. pylori* isolates leads to altered gastric tissue and human protein binding that correlated with modification of the LPS and genetic changes in fucose and galactose transferases. This work highlights LPS as a genetically modifiable adherence factor under selective pressure during stomach colonization.

## Results

### BASEHIT reveals diverse binding phenotypes amongst *Helicobacter pylori* isolates which correlate with genetic phylogeny

To evaluate *H. pylori* binding characteristics, we tested a panel of 48 *H. pylori* strains and isolates for adherence to human exoproteins using the BASEHIT yeast expression library (**39**). Our *H. pylori* panel (**Supplemental Table 1**) included 42 isolates from the J99 culture collection, isolates taken from a single chronically infected subject at two timepoints from multiple biopsy sites (**4, 8, 10**), reference strains J99, PMSS1 (**40**), G27 (**41**), and 26695 (**42**), and mouse stomach-colonizing descendants of these reference strains (**13, 43**). Profiling all 48 *H. pylori* strains in parallel, we quantified yeast barcode counts for the BASEHIT library before and after selections with biotin-labeled *H. pylori* complexed with the human-protein expressing yeast. We then transformed barcode counts into zero-inflated negative binomial interaction (ixn) scores (**Supplemental Tables 2-3**) using the basehitmodel R pipeline (https://github.com/andrewGhazi/basehitmodel) (**39**). In addition to computing ixn scores, the basehitmodel R pipeline annotates these as hits by either meeting the standard or stringent binding thresholds (95% interval excludes zero, estimated effect size > 0.5, and a concordance between replicates of > 0.75 or > 0.95, respectively) determined by Sonnert *et al.* (**39**).

Of the 3324 human proteins represented in the BASEHIT library, 92 bound to one or more *H. pylori* strains (**Supplemental Table 4**). The ixn score data for these 92 exoproteins were used for all subsequent analyses. Specific strains showed enrichment for human exoprotein binding, with SS1 (16 hits), 26695 (13 hits), and J99 culture collection isolates SC8 and D3 (10 hits each) having the most. Some strains of *H. pylori*, including J99, had no hits even though their ixn scores revealed differential interactions within the set of 92 exoproteins. To facilitate comparison of relative binding between strains, we normalized ixn scores for each of the 92 human exoproteins within a given strain by subtracting the minimum ixn score per strain from each then dividing by the difference between the maximum and minimum (**Supplemental Table 3**). This allowed clustering and principal component analysis (PCA) to explore whether human exoprotein binding pattern correlated with genetic relatedness of strains (**Figure 1A-B**). Hierarchical clustering revealed that the binding phenotypes grouped strains according to phylogeny (**Figure 1A** and **Supplemental Figure 1**). Even though J99 had no binding calls, it grouped with the isolates from its same host using our normalization approach. PCA further showed that the most distantly related strains cluster further apart (**Figure 1B**). J99 culture collection isolates all grouped in the PCA and separated from the strains derived from unrelated individuals. Within the J99 culture collection, distinct phylogenetic subgroups described by Jackson *et al*. (**8**) generally clustered together and apart from the unrelated reference strains. Phylogenetic subgroup 2B appears to split between individual isolates being closer to subgroup 2A or subgroup 1A. As orthogonal validation of the lineage-ixn score relationship, we calculated the correlation between average nucleotide identity (ANI) and interaction score using vectorized distance matrices and found it to be moderately positive (Pearson correlation coefficient: 0.45) and significant (p-value: 2.75·10^-55^). ANOVA analyses for each protein, which tested for differences in binding by genetic lineage, found 33 proteins significantly different across lineages (Bonferroni-corrected p < 0.000526), with the 10 lowest p-values belonging to RNASE8, CD99, C19orf18, T4S19, IL17BR, GPR17, INSL4, FMR1NB, OAS1, and MCHR1.

**Figure 1:**
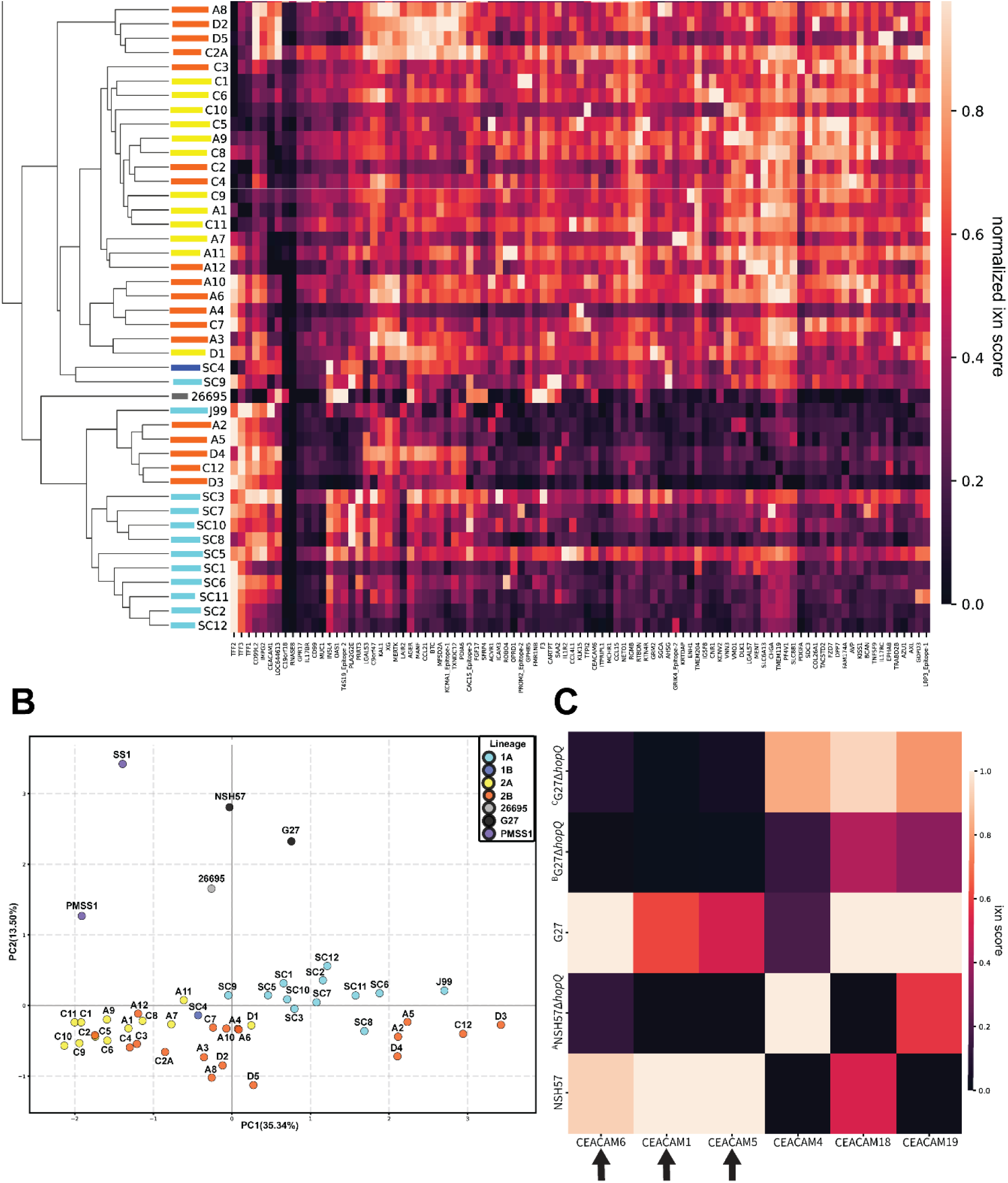
BASEHIT elucidates strain-specific binding of bacteria to human exoproteins. (A) Heatmap and clustergram of normalized basehitmodel (**39**) ixn scores (relative binding) for 92 human proteins that showed significant binding to at least one *H. pylori* strain. Phylogenetic lineages denoted by color (J99 culture collection, labeled according to previously described (**8**) phylogenetic subgroups 1A (cyan), 1B (blue), 2A (yellow), 2B (orange), as well as the several reference strains of *H. pylori* (and their mouse adapted derivatives) PMSS1 and SS1 (purple), G27 and NSH57 (black), C2 and C2A (orange), and 26695 (grey). (B) PCA of *H. pylori* strains (colored by subgroup) based on normalized ixn score. PC1 and PC2 shown (48.84% of total variance). (C) A heatmap showing ixn scores of G27, NSH57, and three isogenic *hopQ* deletion strains ( ^A^ NSH57 *hopQ::*CAT (**45**), ^B^G27 *hopQ::*CAT (**45**), and ^C^G27 *hopQ::aphA3-sacB* (**21**)) for the human CEACAMs present in the 92-protein set. Black arrows indicate CEACAMs showing reduced binding of the mutant compared to the parent wild type strain.

Considering all standard and stringent binding calls, some human exoproteins, such as trefoil factors TFF2 and TFF3, and CD99L2 (CD99 molecule-like two putative adhesion protein) showed binding to many *H. pylori* strains (15, 12, and 11 hits) while most human proteins (65 of 92) were significantly bound by a single strain. In total, there were 186 *H. pylori-*human protein interactions identified. Of these, 33 were stringent binding interactions, with the trefoil factors TFF2 and TFF3 enriched with 10 and 6 hits, respectively. These proteins have been previously described to bind to *H. pylori* LPS (**23, 25, 26, 44**).

As an internal validation, we included a set of isogenic *hopQ* (*omp27*) mutants of strain G27 and its mouse-adapted derivative NSH57 (**Figure 1C**). Despite G27 and NSH57 having no basehitmodel standard or stringent binding calls to any CEACAMs, analysis of ixn scores revealed reduced binding to CEACAM 1, CEACAM 5, and CEACAM 6 upon *hopQ* disruption, consistent with previous characterizations of HopQ as a CEACAM-targeting adhesin (**19**). Collectively, these data indicate that BASEHIT-derived ixn scores provide a useful measure of *H. pylori* strain differential binding of human exoproteins and that strains have unique human exoprotein binding signatures that correlate with genetic phylogeny.

### Genetic variations associated with human protein binding show enrichment for bacterial cell envelope-related genes

To identify the genetic causes of differential binding, we performed two genome-wide association studies (GWASs) using the bugwas (https://github.com/sgearle/bugwas) (**46**) and pyseer (https://github.com/mgalardini/pyseer) (**47**) pipelines on the 92 human proteins with binding phenotypes against SNPs and InDels in the J99 culture collection as covariates (**Figure 2**). These GWAS pipelines were chosen for their lineage or locus-based correction, necessary given *H. pylori’s* high degree of genetic diversification and the uneven sampling of genetic lineages in our collection. Based on a p-value cutoff of 0.0001, 60 polymorphisms showed significant associations with binding phenotypes across 35 distinct loci in the combined GWAS dataset. Most of the polymorphisms (35/60) mapped to loci related to the bacterial cell surface (**Table 1** and **Supplemental Table 5**). Of these 35, seven mapped to intergenic regions near OMPs, three of which are likely to drive promoter phase variation since they reside in homopolymeric tracts expected to alter spacing between the -35 and -10 promoter elements. We also identified 22 polymorphisms within the coding region of OMPs associated with binding phenotypes, including uncharacterized putative OMPs, as well as OMP loci previously linked to virulence (*sabA, sabB,* and *babB*) with significant GWAS p-values. The remaining cell surface-related hits mapped to loci encoding proteins that build structural components of the cell wall, including the cell wall polymerase PBP-1, and the enzyme *gmhA*, which synthesizes phosphoheptose used in LPS biosynthesis, as well as polymorphisms in *csd4* and *csd5*, helical cell shape-determining genes. Several human proteins showed significant associations with multiple *H. pylori* genes, with PLA2G2E (21/60) having the most. Many of the human proteins have known glycosylation, such as KLK15 (**48**), VNN3 (**49**), and TFPI2 (**50**). Thus, the *H. pylori* adhesins previously shown to bind specific glycan epitopes that show binding associations with these proteins such as SabA, SabB, and BabB, may interact with their glycans rather than the polypeptides.

**Figure 2:**
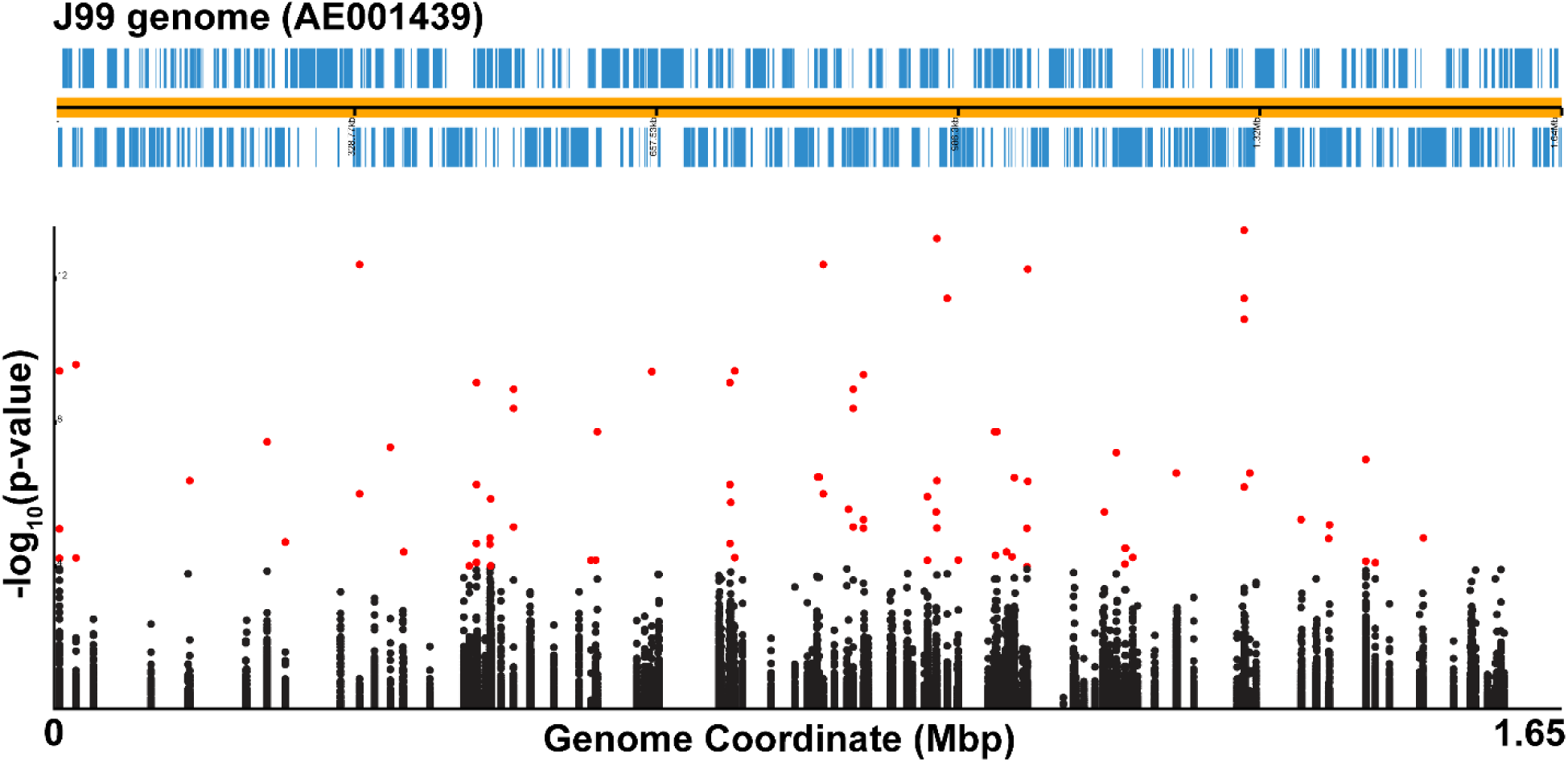
GWAS of SNPs/InDels and binding reveals potential genetic associations. A Manhattan plot of combined data from two GWAS analyses (bugwas (**46**) and pyseer (**47**)) of basehitmodel ixn scores associated with J99 culture SNPs and InDels, aligned to the J99 genome (Accession: AE001439). To reduce noise associated with small sample size and genetic relatedness, repetitive p-values were eliminated. A p-value cutoff of 0.05 was applied for inclusion in the graph, and a more stringent 0.0001 (-log_10_(p-value) ≥ 4) cutoff (red dots) was used to determine significance.

**Table 1:** Differential binding of human exoproteins is related to polymorphisms at discrete loci.

| Human Protein<sup>a</sup> | *Hp* loci [-log(p-value)]<sup>b</sup> |
| --- | --- |
| C9orf47 | JHP0928 [4.34] |
| CAC1S | *homB* (2 SNPs) [5.10] |
| GDPD3 | JHP0305 [12.50] *gmhA* [8.46] |
| GRIK4 | *sabA* [4.27] JHP0953 [4.02] JHP0953 (2 SNPs) [4.02] JHP0955 [6.41] JHP1053 [4.54] |
| IL1R2 | JHP1300 [4.17] |
| KLK15 | *sabB* [5.82] *csd5* [6.64] |
| KRTDAP | *sabA* [9.51] JHP0955 [12.37] JHP1272 [4.81] |
| LRP3 | JHP0305 [6.06] *gmhA* [5.13] |
| MERTK | JHP0440 [4.64] |
| OAS1 | *atpG* [4.28] |
| PDGFA | JHP0589 [9.49] *hopK* [5.98] *hopQ* (4 SNPs) [6.95] |
| PLA2G2E | PBP-1A [4.20] *hsdM* [5.63] *homB* (2 SNPs) [13.23] JHP0928 (2 SNPs) [7.81] *babB* (3 SNPs) [11.55] *babB* [13.46] JHP1300 [7.03] JHP1272 [5.19] JHP1312 [4.13] |
| SGCA | *csd4* [4.43] *hopK* [4.20] JHP0937 [4.43] |
| SLC8B1 | *pepF* [4.04] |
| TFPI2 | JHP0440 [4.04] *sabB* [9.18] JHP0757 [6.53] |
| TRABD2B | *deaD* [4.71] |
| VMO1 | *sabB* [4.67] |
| VNN3 | *sabB* [6.32] |

<sup>a</sup>Human proteins with significant binding phenotype hits in the GWASs described in **Figure 2**.
<sup>b</sup>*H. plori* loci with coding region SNPs or InDels that met the filter thresholds of non-repetitive p-values < 0.0001. For information about the mutation context and location see **Supplemental Table 5**.

### Mouse-adapted *H. pylori* isolates show increased stomach tissue binding associated with altered LPS O-antigen profiles

In addition to the J99 culture collection isolates, we further evaluated within-host evolution in our panel through the inclusion of three pairs of strains in our analysis that had undergone passage through mice to create mouse stomach colonizing strains (PMSS1 and SS1, G27 and NSH57, and C2 and C2 adapted – **C2A in figures**). For all three pairs, the post-mouse isolates displayed dramatically altered ixn scores that included both gain and loss, but on balance, higher overall human protein binding than the cognate parent strain (**Figure 3A**). While all three pairs exhibited higher average ixn scores across the 92 proteins, only NSH57 had statistical significance compared to its parent, G27 (p-value = 0.027, Mann-Whitney U). SS1 had 16 binding calls while PMSS1 had four, of which three overlap with SS1 (**Supplemental Table 4**). Neither the parent C2 isolate, nor C2 adapted had hits passing either threshold using basehitmodel. To assess the biological relevance of these profound changes in human protein interactions, we explored whether these strains showed altered adherence to *ex vivo* mouse stomach tissue. As shown in **Figure 3B**, in all three cases, the mouse-adapted isolates (SS1, NSH57, and C2 adapted) had higher adherence to mouse gastric tissue than their pre-mouse parent strain (PMSS1, G27, and C2).

**Figure 3.**
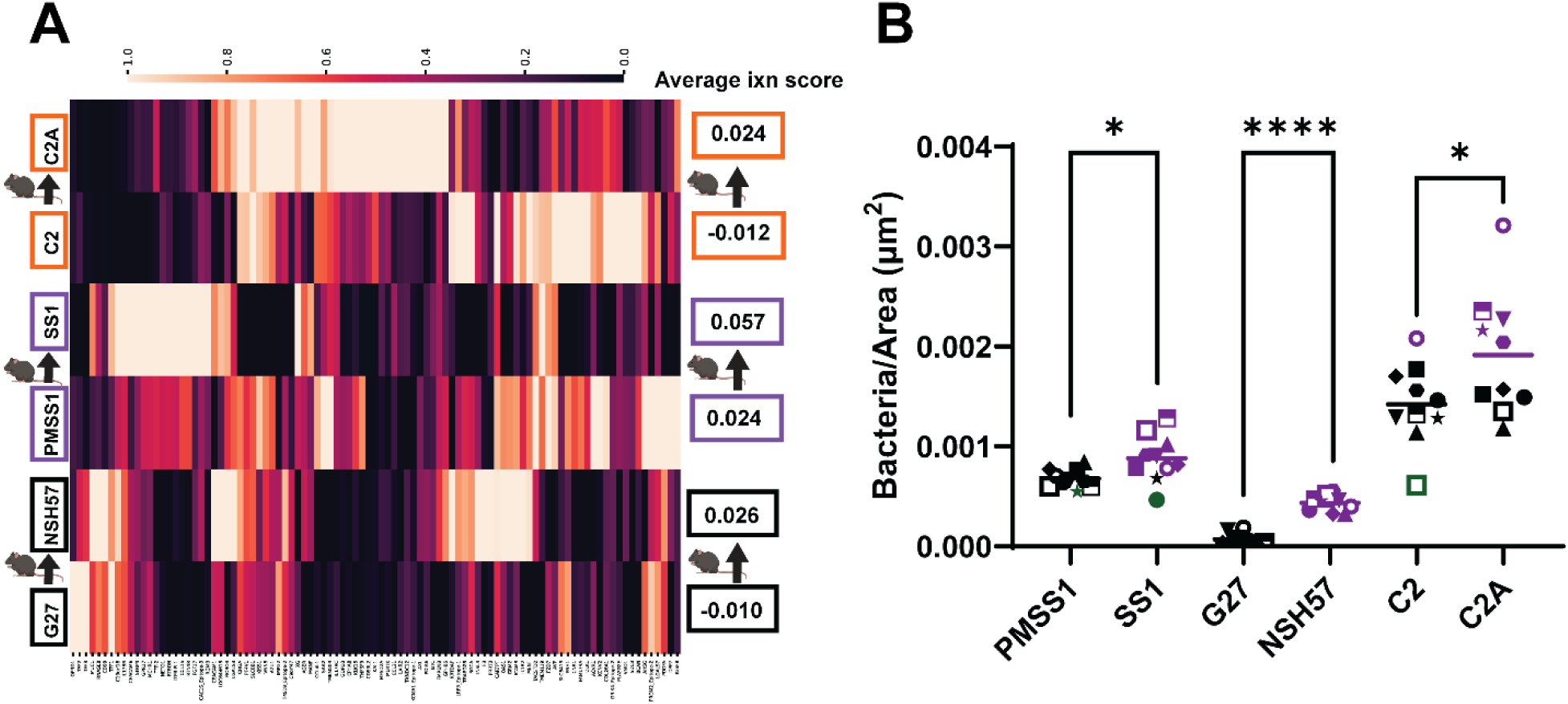
Mouse-adapted strains have altered human exoprotein binding patterns and bind mouse stomach tissue with greater affinity. (A) A heatmap of ixn scores from G27, PMSS1, and C2, as well as their cognate post-mouse stomach infection isolates (NSH57, SS1, and C2A). Average ixn score across 92 *H. pylori*-binding target proteins displayed. (B) Adherence of fluorescein-labeled bacteria of the indicated strain/genotype to FFPE C57BL/6NJ mouse stomach tissue slices. Number of adhered bacteria per area (µm^2^) of tissue plotted. Purple - values greater than one standard deviation above wild type average, green - values one standard deviation below. Sequential tissue slices from the same animal are graphed with the same symbol. Statistical significance determined by paired T-test in GraphPad Prism, *p-value < 0.05, ****p-value < 0.0001.

Given that we saw binding differences to the trefoil factors that are known to bind LPS, we tested whether LPS modification could underlie the differences in adherence by our mouse-adapted isolates. To test changes in lipid A, which could affect presentation of the LPS O-antigen attached to it, we performed Fast Lipid Analysis Technique (FLAT) which selectively cleaves the proximal Kdo (3-deoxy-d-manno-octulsonic acid) from lipid A allowing for the free lipid to be analyzed on-surface with MALDI mass spectrometry (**51**). None of the three pre- and post–mouse-adapted strain pairs showed any detectable alterations in lipid A (**Supplemental Figures 2-4**).

Direct modification of the LPS core oligosaccharide and/or O-antigen could also occur through genetic modification and phase variation of LPS glycosyltransferases. To test this possibility, we extracted the LPS from PMSS1, SS1, G27, NSH57, C2, and C2-adapted and analyzed the electrophoretic mobility of the LPS oligosaccharide qualitatively via SDS-PAGE. Each of the post-mouse adapted isolates harbored a mobility shift in the band corresponding to the complete LPS chain with a modal species of O-antigen repeat length (**Figure 4A**). The magnitude of the electrophoretic shift varied by strain background, with PMSS1 and SS1 showing the largest changes in mobility, but in all cases the altered patterns were consistent with LPS modifications that increased electrophoretic mobility after mouse passage. These band shifts suggest structural modifications to the LPS oligosaccharide chain that either reduce the modal number of repeated glycan units to make the molecule shorter or alter oligosaccharide composition to affect charge. To explore the genetic basis for altered LPS structure, we utilized existing whole genome sequences (WGS) of PMSS1 (GenBank: CP018823.1), SS1 (GenBank: CP009259.1), G27 (GenBank: CP001173.1), C2 (BioProject: PRJNA622860), and C2-adapted (GenBank: CP089284.1). We performed Oxford Nanopore long read sequencing (Plasmidsaurus) for NSH57 (**Supplemental Table 7**). As noted previously (**6**), SS1 has phase variation in the c-terminus of *futB,* which encodes a FucT α-(1,3)-fucosyltransferase and in the promoter of *rfaJ* (Lewis antigen presentation-associated α-(1,2)-glucosyltransferase) compared to PMSS1 (**Figure 4F**). Our sequencing data revealed that NSH57 carries a frameshift that restores *HpG27_0579/80* to a single ORF encoding a *HPYLPMSS1_0826* β-(1,3)-galactose transferase homolog, which was not included in the recent redescription of G27 LPS synthesis (**30–32, 52**). NSH57 also displays *futB* (*HpG27_0613*) repeat contraction and early termination of *futC* (*HpG27_0086*). C2 adapted harbors a single non-synonymous SNP in *sabB.* However, many LPS genes show phase variation in simple repeats (**33–35, 53**) that all WGS platforms struggle to accurately measure. Indeed, the original sequencing of SS1, when compared to PMSS1, missed the changes in these relevant loci that were later defined using additional shotgun sequencing (**6**).

**Figure 4.**
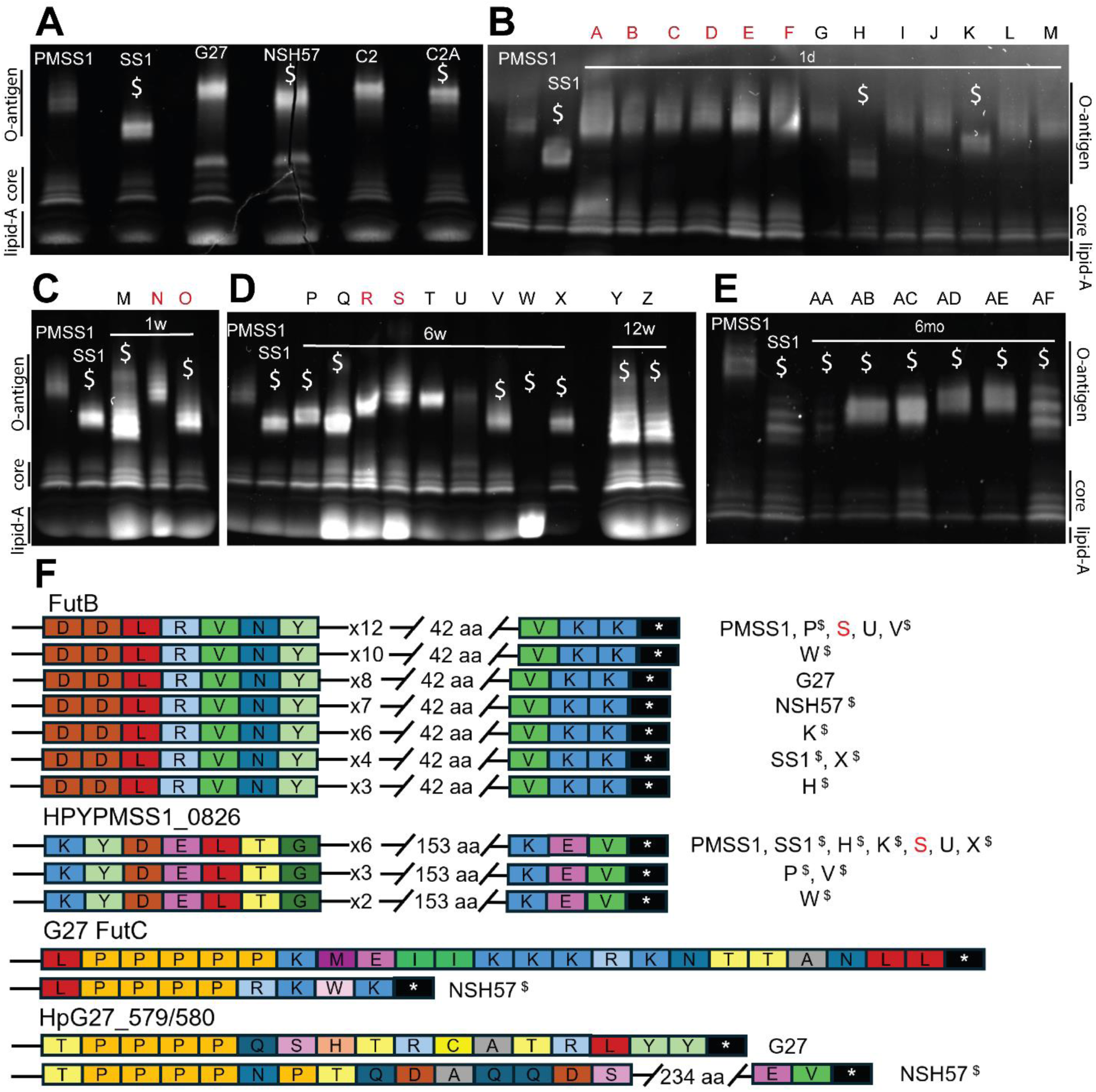
LPS structural variation and glycosyltransferase genetic variation occurs frequently during mouse infection. (A-E) SDS-PAGE of purified LPS stained with ProQ Emerald 300 kit. Higher mobility LPS denoted ($). (A) Previously described *H. pylori* mouse-adapted strains and their parent human isolate. (B-E) PMSS1, SS1, and mouse outputs of PMSS1 at indicated times post-infection in C57BL6/N (black) or GIM mice (red). (F) Predicted amino acid variants of the α-(1,3)-fucosyltransferase FutB, the putative galactosyltransferase HPYLPMSS1_0826, G27 FutC (HpG27_0086), and the HpG27_579/580 loci from the subset of strains we sequenced. Strains and isolates are indicated next to the allele they are predicted to encode based on WGS.

To assess the frequency and timing of LPS structural alterations during mouse passage of human isolates, we screened a set of mouse outputs from multiple independent experimental infections of PMSS1 in either the normal, wild type stomach (C57BL/6NJ) or a mouse model of gastric intestinal metaplasia (GIM) that was used in the mouse adaptation of C2 adapted (**13**). The GIM model uses transgenic expression of an oncogenic allele of *Kras* (*Kras^G12D^*) in chief and a subset of gastric epithelial precursor cells for 6 weeks to induce GIM (**20, 54**). As shown in **Figure 4 B-E** (and **Supplemental Figure 5A-D**), we observed LPS gel mobility alterations in as short as one day of mouse infection. As with the previous isolates tested, several different magnitudes of electrophoretic shift were observed, despite all of these outputs being derived from PMSS1. Nearly all of them represented higher mobility O-antigen modal moieties. In output isolates from wild type C57BL6/NJ mice, we observed electrophoretic shifts (to higher mobility species) in a subset of one-day (5/33), one-week (2/2), six-week (5/7), 12-week (2/2), and six-month (15/16) outputs tested. In the GIM model, fewer outputs showed altered LPS mobility (0/6 one-day, 1/2 one-week, and 0/2 six-week) than in isolates from wild type C57BL/6NJ mice. As with the more characterized mouse-colonizing and parental strain pairs, we used FLAT to screen for lipid A modifications in four PMSS1-derived mouse outputs (Isolates M, P, O and Z in **Figure 4C**). We detected a 43 *m/z* decrease, consistent with substitution of the canonical *H. pylori* ethanolamine (123 *m/z*) with a phosphate group (80 *m/z*) on the lipid A glucosamine (**55–57**) in Isolate P (**Supplemental Figure 6**), a single 6-week mouse infection derivative of PMSS1, which we then validated with MS/MS on both PMSS1 and Isolate P to derive the structures (**Supplemental Figures 7 and 8**). This was the only lipid A modification observed in our dataset. Collectively, these results reveal that LPS structural modifications that result in electrophoretic mobility shifts are frequent during mouse infection and can occur early in infection, suggesting a potential role during initial establishment of colonization that continues to be favored during persistent infection. However, these LPS mobility changes do not appear to reflect changes in lipid A structure.

### Alteration of LPS O-antigen composition modulates gastric tissue adherence

Based on our observation of adherence correlated LPS structural changes, we evaluated the loss of function of individual glycosyltransferases from the recently recharacterized *H. pylori* G27 LPS biosynthetic pathway. A comprehensive list of glycosyltransferases discussed is available in **Supplemental Table 6**, adapted from Tang *et al*. (**58**). Several mutations completely truncate the oligosaccharide so that no O-antigen is present, which differs from the electrophoretic shift observed in the mouse outputs; however, the *futB* mutant displayed a modest increase in electrophoretic mobility similar to NSH57 (**Figure 5A**). We evaluated each deletion mutant in the mouse tissue adherence assay used in **Figure 3B**. We found that isogenic mutants of G27 lacking O-antigen had similar or lower adherence to their parent strain (**Figure 5C**). However, deletion of *HpG27_0761* and *HpG27_1236*, which results in loss of the proximal D,D-manno-heptose and/or its sidechain but still allows ligation of O-antigen to the second L,D-heptose of the core and isometric LPS (**Figure 5A-B**) (**32, 52, 59**), led to increased adherence compared to wild type G27. These data indicate that modulation of the O-antigen or the core oligosaccharide sidechain of *H. pylori* LPS modifies gastric tissue adherence properties, but none of the deletion mutants, except for the deletion of *futB,* showed LPS mobility changes that matched those observed in the mouse output strains, and the *futB* deletion strain did not show altered tissue adherence.

**Figure 5.**
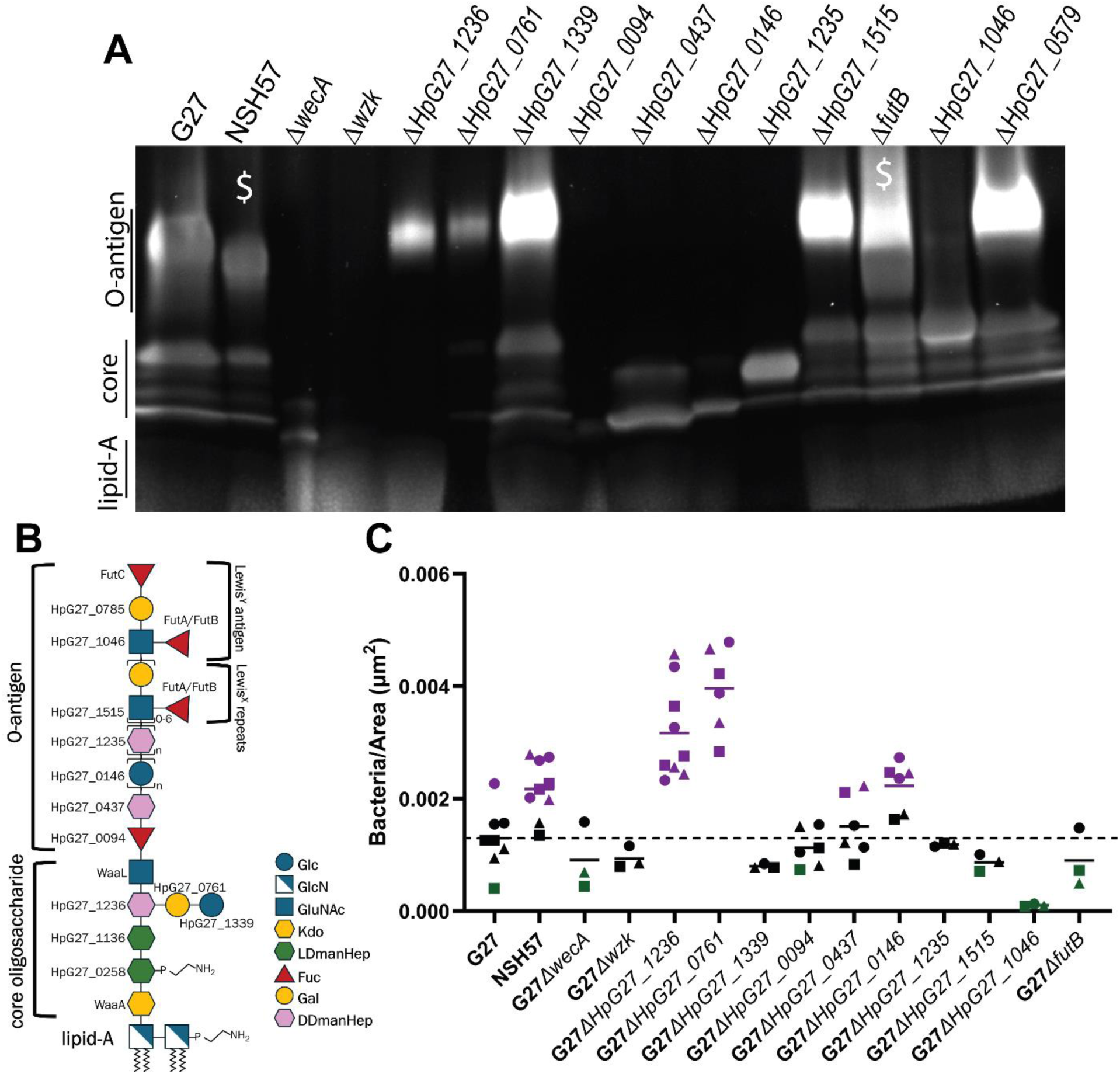
Deletion of specific glycotransferases increases adherence to mouse stomach tissue. (A) SDS-PAGE of purified LPS from a panel of deletions subset of the characterized G27 glycosyltransferases. Loss of O-antigen production was identified in *wecA, wzk, HpG27_1046, HpG27_0437, HpG27_0094,* and *HpG27_1235* deletions. (B) Schematic of the G27 LPS structure with enzymes responsible for the addition of each unit, adapted from Li et al. (**32**). (C) Adherence of fluorescein-labeled *H. pylori* to FFPE GIM mouse stomach tissue. Number of adhered bacteria per area of tissue plotted in GraphPad Prism. Purple - values greater than one standard deviation above wild type average (dotted line), green - values one standard deviation below. Sequential tissue slices from the same animal graphed with the same symbol.

### LPS structural variation correlates with distinct genetic polymorphisms and alters mouse colonization

To directly assess changes that underlie the observed LPS mobility shifts that arise during mouse stomach passage, we sequenced a subset of the PMSS1 mouse outputs (two 1-day and four 6-week) with Oxford Nanopore (Plasmidsaurus). We observed frequent, similar mutations in *HPYLPMSS1_0826* (3/6) and *futB* (4/6). *HPYLPMSS1_0826* encodes a putative glycosyltransferase family 25 galactose transferase and is a homolog *HpG27_0579/580*, which is altered in the NSH57, and a member of the same family as *HpG27_0761*, which has an adherence phenotype when knocked out. Both *HPYLPMSS1_0826* and *futB* contain simple heptameric amino acid repeats in the C-terminus, which shift in number between isolates due to phase variation (**Figure 4F**). All isolates with LPS mobility shifts had repeat length shortening of one or both loci (**Supplemental Tables 9-16**). Additional polymorphisms were found in *HPYLPMSS1_1046* and in the promoters of *HPYLPMSS1_1047* and *rfaJ.* Isolate P, which exhibited altered LPS mobility and carried phosphate rather than phosphoethanolamine on its lipid A glucosamine, did not have any detectable polymorphisms in the phosphoethanolamine transferase EptA. It instead harbored an R159H substitution in LpxE (**Supplemental Table 11**), the 1-phosphatase which catalyzes the first step of the constitutive *H. pylori* lipid A modification pathway (**60, 61**). This *lpxE* polymorphism may cause this altered lipid A phenotype, as *lpxE* inactivation has been demonstrated previously to result in similar shifts to phosphate moieties by disruption of ethanolamine addition (**61**). Loci not related to LPS synthesis also differed in all sequenced isolates, although many of these polymorphisms were detected in the resequencing of our lab’s stock of PMSS1 (**Supplemental Table 8**), suggesting some stochastic variation in sequences due to *in vitro* growth. Among the one-day mouse outputs with shortened LPS (H and K in **Figure 4C**), which are expected to have fewer polymorphisms than in the longer infections, both had *futB* c-terminal repeat length changes. One isolate (H) also had additional SNPs in *futB,* while the other (K) had *rfaJ* promoter variation. We also sequenced two isometric LPS isolates from the six-week outputs (S and U in **Figure 4C**) and found that they did not contain polymorphisms in any of these loci (**Supplemental Tables 12-13**). Our genetic analyses indicate that multiple loci may be responsible for *in vivo* structural modification of LPS. Changes in either *futB* or *HPYLPMSS1_0826* seem to be necessary, with modifications of just one being sufficient to alter LPS gel mobility.

Given higher frequency of *futB* polymorphisms and heptad repeat loss as soon as one day post-infection, we focused on the behavior of an isolate with futB shortening during competitive infection. We coinfected C57BL/6NJ mice with an equal mixture of PMSS1 and either the isometric one-day output strain (G or I in **Figure 4C** and **Figure S8**) or an increased LPS electrophoretic mobility one-day output strain that had the shortest number of heptad repeats (three) in *futB* and full-length heptad repeats in HPYPMSS1_0826 (H in **Figure 4C**). To distinguish among the input strains in replicate experiments, we introduced a kanamycin resistance cassette at a neutral locus in either PMSS1 or the output strain, using both male and female mice. The colonization and competition phenotypes were variable across three biological replicates and did not correlate with sex or strain which carried the antibiotic resistance cassette. One week after infection, the isometric isolates (G and I) either modestly outcompeted or were modestly outcompeted by PMSS1, depending on the mouse and replicate (**Figure 6A-B**). The higher LPS gel mobility strain (H) had a large competitive advantage in one replicate (eliminating PMSS1 in one mouse), but a more modest advantage and disadvantage across the other two replicates (**Figure 6C**).

**Figure 6.**
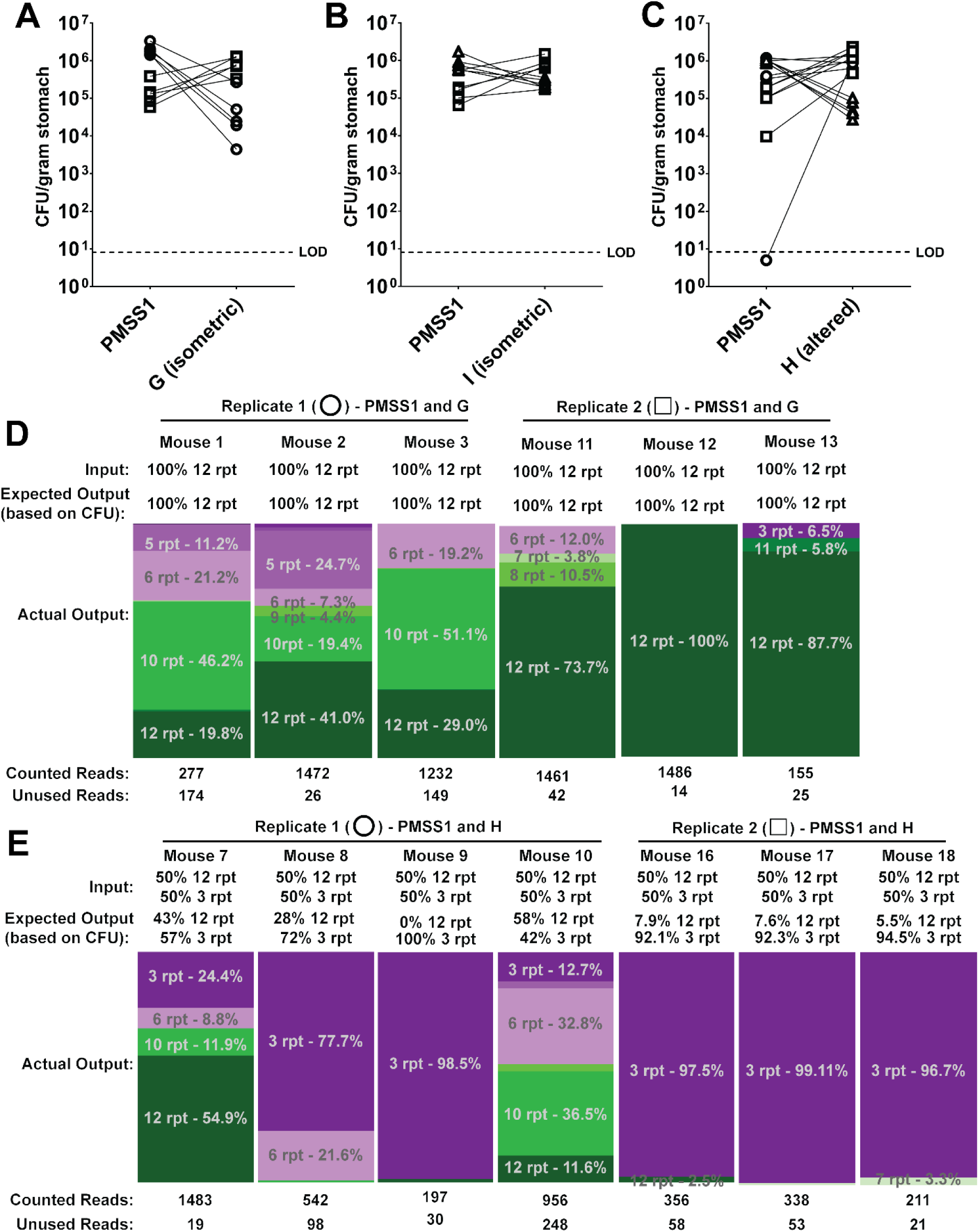
Coinfection of mice with PMSS1 and single day outputs reveals colonization differences and additional *futB* selection. (A-C) Colonization data of PMSS1 and either an (A-B) isometric one-day isolate (G or I in Figure 4C) or an (C) altered LPS electrophoretic mobility one-day isolate (H in Figure 4C) coinfected into 10 C57BL6/N mice each for 1 week. Points represent the recovered CFU per gram of stomach tissue from each mouse and lines connect output strains from the same mice. LOD: limit of detection. (A-C) Points with the same shape indicate data from the same biological replicate. Replicates 1 (open circle) and 3 (open triangle) were performed with cassette-marked Isolates G, I, or H in female mice, while Replicate 2 (open square) was performed with cassette-marked PMSS1 in male mice. (D-E) Amplicon sequencing of *futB* from the stomach homogenate of representative mice from (D) PMSS1/G or I or (E) PMSS1/H coinfections displayed as the percentage of sequencing reads harboring each repeat length of the C-terminal 21-bp repeat (rpt) motif. Input and expected outputs are based on WGS and CFU data.

Variation in *futB* appeared most often in our altered LPS isolate WGS data and was the context in which Isolate H was altered. Therefore, we evaluated new *futB* C-terminal repeat contraction occurring during coinfection, which may contribute to the variability in colonization of individual isolates. We extracted DNA from homogenized stomachs used for CFU plating and used amplicon sequencing of the *futB* locus of three to four mice per condition per replicate, counting the number of 21 base pair repeats present in each read. Of the 22 stomach homogenates sequenced, 20 had a reduction in the number of PMSS1-like 12-repeat isoforms detected in the output compared to the input (**Figure 6D-E** and **Supplemental Figure 9A-B**).

Conversely, 5 of 10 mice infected with the mix containing three repeat isoforms had increases in the three-repeat length in their output. Other repeat lengths, not present in the inoculum were also detected in 18 of the 22 mouse stomach homogenates, with repeat lengths of five, six, seven, and 10 appearing most often alongside the input repeat lengths of 12 and three. In 15 of the 22 mice, the proportion of 12-repeat isoforms was less than that of all shorter repeat motifs. We also applied our amplicon sequencing strategy to liquid-culture grown PMSS1 and Isolates G, H, I, J, and K, which replicated the WGS data and demonstrated that there were no detectable mixed populations in input PMSS1, G, H, and I used in the infections at our detection limit (approximately 10^-3^, **Supplemental Figure 9C)**. These data suggest that LPS structural changes likely affect colonization in addition to adherence and that *futB* changes are dynamic during infection.

## Discussion

In this study, we utilized the BASEHIT yeast expression library to profile the interactions between diverse *H. pylori* strains and a comprehensive library of host proteins, revealing lineage specific interactions between *H. pylori* and human protein epitopes. Interestingly, some *H. pylori* strains (SS1, 26695, SC8, and D3) exhibited a broad binding phenotype similar to the “superbinders” identified by Sonnert *et al.* (**39**) in their debut of BASEHIT; while others, such as the reference strain J99, had no significant binding at all. Heterogeneous binding by *H. pylori* is unsurprising, given the species’ relatively high genetic diversity; however, even *H. pylori* isolated from a single human host exhibited broad protein interaction differences akin to what was seen in a multiple phyla dataset for Sonnert *et al.* (**39**). Our GWASs also identified targets for future study on both the human and the bacterial side. Most of these *H. pylori* loci are related to cell surface modifying genes and their promoters. Known lectin-like adhesins, SabA and BabA, had significant hits for binding to multiple proteins. While BASEHIT is intended to elucidate interactions with proteins, post-translational modifications like glycosylation are still possible, albeit with the yeast glycosyltransferases rather than the human glycosylation machinery. It is unclear whether the different variants of these proteins are directly modulating their ability to bind to host proteins or protein glycans.

We identified a pattern of modified binding after mouse stomach passage by distinct *H. pylori* strains even though BASEHIT characterizes human exoproteome binding. This host species discrepancy may explain protein ixn scores that both increased and decreased, while trending higher overall. We hypothesized that such a broad change in adherence resulted from a broad change in bacterial physiology, rather than variation of a single adhesin. This led us to examine structural modifications of LPS. We identified a modification of lipid A in a single isolate which we hypothesize is caused by a *lpxE* polymorphism reducing its enzymatic efficiency for dephosphorylating lipid A and thereby reducing subsequent addition of phosphoethanolamine. Prior literature suggested that LpxE inactivation and subsequent lipid A phosphate decoration reduced mouse stomach colonization and sensitized the bacteria to polymyxin (**61, 62**), despite our observation of this apparent loss of LpxE activity arising during mouse infection and being isolated from media containing polymyxin B. While intriguing, our altered LPS phenotype more closely correlated with changes in qualitatively determined LPS O-antigen structure, with mouse-adapted LPS migrating faster during gel electrophoresis due to either a reduction in polysaccharide length or a change in the glycan composition of the molecule.

We first examined how alterations in LPS O-antigen structure affect gastric tissue adherence by individually disrupting the glycosyltransferases that comprise the recently redefined complete G27 LPS biosynthetic pathway (**30–32, 58**). We replicated the electrophoretic shift pattern observed in the G27 derivative NSH57 only through genetic disruption of the α-(1,3)-fucosyltransferase, *futB*. G27 and PMSS1 each exhibit different LPS glycosyltransferase compositions, with PMSS1 harboring the East-Asian characteristic lack of *HP1283* and *HP1578*, duplication of *HP1105* (*HPYLPMSS1_1046* and *HPYLPMSS1_1047*), and presence of *JHP0562* (*HPYLPMSS1_0825*) (**32**). This difference in LPS synthesis strategy may explain why loss of *futB* did not have a significant impact on adherence in G27 and why the electrophoretic shift observed in G27 is more subtle. We also recognize that genetic disruption alone may obscure the nuance of phenotypes, especially if there is some functional redundancy by members of the same glycosyltransferase family or if these enzymes have more roles than previously characterized. We did, however, identify several G27 glycosyltransferase loci that do increase adherence when disrupted, *HpG27_0761* (a glycosyltransferase family 25 galactose transferase) and *HpG27_1236.* These mutants exhibited isometric LPS via gel as they affect a sidechain, though they are more similar to our naturally occurring variants in that they do not completely inhibit O-antigen production.

To gain further insight into the LPS changes observed during mouse stomach passage, we sampled PMSS1 mouse infection outputs and correlated *in vivo* LPS structural changes with C-terminal heptad amino acid repeat length changes in FutB and glycosyltransferase family 25 galactose transferases, which, in the case of FutB, is consistent with prior literature on its heptad repeat length variation and function (**33, 37, 38**). We note, however, that detecting LPS gene variation by sequencing is complicated by the highly repetitive nature of these loci, which none of the major sequencing platforms measures accurately. This limitation was partially mitigated in our newly sequenced isolates by using gentle DNA extraction methods that increased fragment size and thereby improved recovery of long reads. We also compared sequencing performed using different platforms, which could bias our conclusions. However, our sequence data consistently suggested variation in the same loci or in members of the same glycosyltransferase family. Polymorphisms in these loci were also detected in the better characterized mouse-colonizing strains SS1 and NSH57 further supporting their importance.

Furthermore, variation in FutB heptad repeat length appeared more frequently and as early as one day post-infection. In competition with the PMSS1 parent strain, mouse output strains with full length or shorter FutB heptad repeat lengths persisted with apparently stochastic colonization advantage that did not depend on sex or whether the parent or output strain carried the antibiotic cassette used to distinguish among the two infection strains. Amplicon sequencing of stomach homogenates revealed that while heptad repeat length appears stable during broth culture to the limit of our detection (∼10^-3^), in all but two mice of 22 mice analyzed the frequency of full length (12 heptad repeats) were lower than predicted by CFU. In addition to the two parental genotypes, we observed a range of heptad repeat lengths and inferring from the CFU data both expansion and contraction of heptad repeats accumulated during infection. The remaining two mice with significant 12 repeat populations may also harbor other polymorphisms, such as *HPYPMSS1_0826* C-terminal repeat variation, which we would expect to confer a colonization advantage as well.

While phase variation in LPS and other cell envelope genes has been extensively documented (**4, 33–35, 53, 63, 64**), these genetic changes likely occur by recombination. The *futB* C-terminal repeat length variation is structurally similar to the recombination-mediated alteration of the *cagY* middle repeat region, which results in diverse in-frame gene lengths under selective pressure by the host immune response to Cag-T4SS-mediated immune insult (**65–67**). This *cagY* variation occurs at high levels *in vivo* (> 80% of wild type mice infected with *H. pylori* for eight weeks (**66**)) due to selection, similar to what we observe in *futB*; however, *cagY* recombination was detected at much lower frequencies (< 7% of clones tested (**67**)) during growth *in vitro.* Likewise, 50 serial passages performed by Nilsson *et al*. (**38**) were unable to experimentally detect *futB* or LPS alteration in strains grown with standard *in vitro* population selection and bottlenecks, only finding variation in populations passaged 50 times through small bottleneck selections. Given the depth of our broth culture amplicon sequencing, our data suggest *futB* recombination occurs at an even lower rate (a frequency < 10^-3^) than *cagY* recombination events *in vitro* and may possibly arise *de novo* in the stomach.

Based on gel appearances, we hypothesize that the LPS structural change results from alteration of the modal repeat length of the O-antigen Lewis units. The base N-acetyllactosamine (LacNAc) units are synthesized by the addition of galactose to N-acetylglucosamine (GlcNAc) by glycosyltransferase family 25 enzymes (**68**). The LacNAc is then fucosylated by FutB (acting as a heterodimer with FutA) to make the Lewis^X^ (Le^X^) repeats, which are added to the growing LPS chain as a distal homopolymer (**32, 37, 69**). At a lower frequency, the terminal Le^X^ is further fucosylated by FutC to generate Lewis^Y^ (Le^Y^), capping the O-antigen chain and preventing more units from being added (**32, 36**). We propose a model (**Figure** 7) in which LPS varies in length due to the competition between these enzymes for substrate and their combined competition with the terminal fucose capping enzyme FutC. The C-terminal repeat region of FutB occurs in the domain that forms a leucine zipper with FutA and is predicted to be important for proper dimerization and substrate specificity (**69–71**). Changes in repeat length affect the distance between the N-terminal catalytic domain and the distal C-terminal membrane-associated hydrophobic domain. Alterations in this distance, the dimer strength with FutA, or the specificity of substrate could all influence the enzymatic efficiency of the FutA/B dimer and result in changes in the frequency at which FutC beats the addition of the next Le^X^ and terminates the chain, causing short oligosaccharide chains with lower molecular weight and higher electrophoretic mobility. We found that reduced FutB C-terminal repeat length is selected *in vivo,* improving adherence to mouse tissue and colonization in a mouse model. Specific modalities of repeat length (3, 6, and 10 repeats) consistently emerged during our mouse infections, suggesting optimal repeat lengths for production of LPS while in the stomach. We think these collective data suggest that *futB* repeat contraction results in shorter LPS chains, which is favorable *in vivo* and increases adherence.

**Figure 7:**
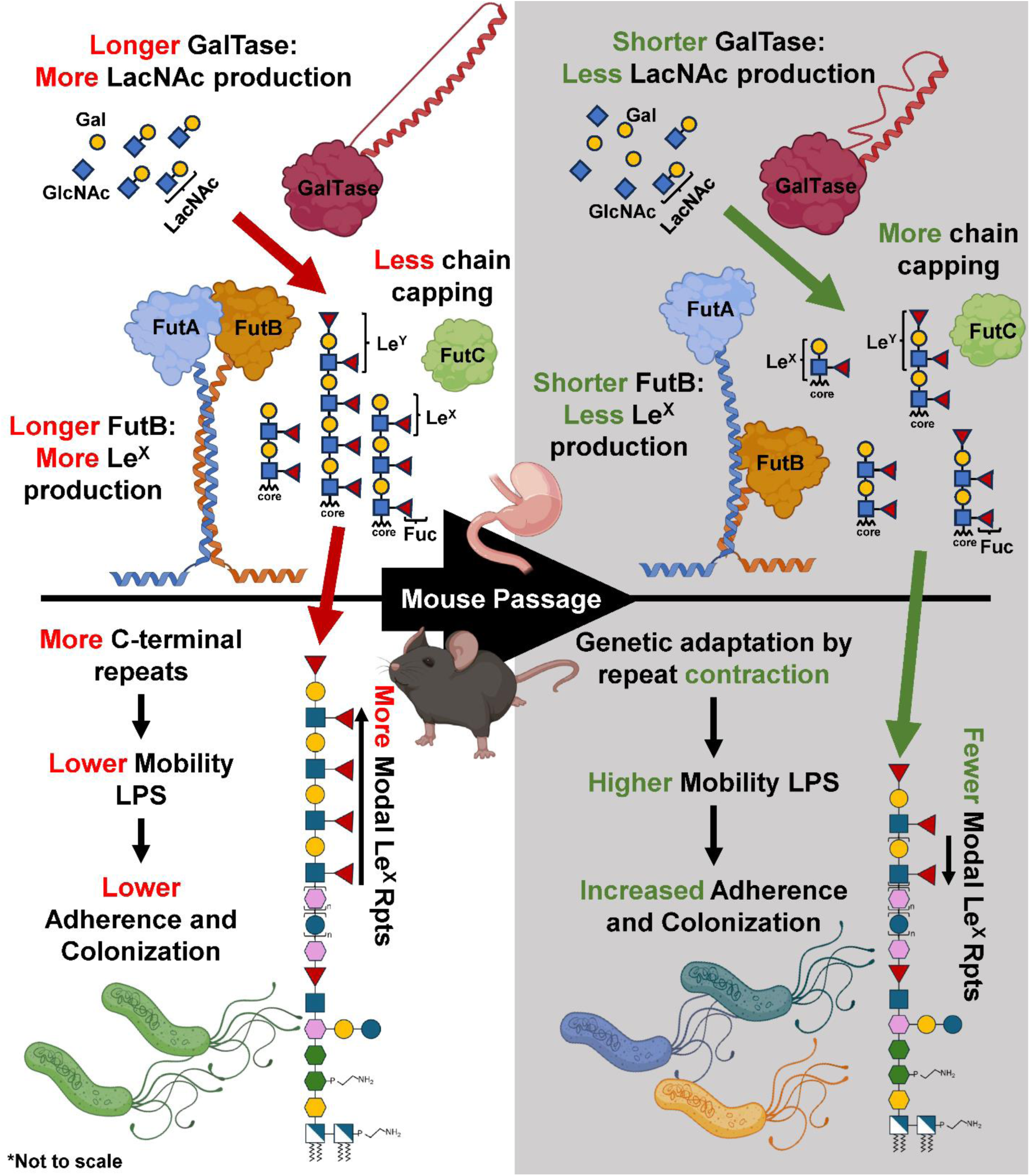
A model of *H. pylori* LPS O-antigen alteration by structural variation of enzymes in competitive reactions. Cartoon representations of galactose (Gal) and fucose (Fuc) transferase (Tase) activity, synthesizing the base N-acetyllactosamine (LacNAc) and downstream Lewis^X^ (Le^X^) repeats and Lewis^Y^ (Le^Y^) caps, before and after genetic adaptation during mouse stomach colonization. Created in BioRender. Frick, J. (2026) https://BioRender.com/e113ia5

## Materials and Methods

### Ethics statement

All mouse experiments were performed in accordance with the recommendations in the National Institutes of Health Guide for the Care and Use of Laboratory Animals. The Fred Hutchinson Cancer Center is fully accredited by the Association for Assessment and Accreditation of Laboratory Animal Care and complies with the United States Department of Agriculture, Public Health Service, Washington State, and local area animal welfare regulations. Experiments were approved by the Fred Hutch Institutional Animal Care and Use Committee, protocol number 1531.

### H. pylori culture

The *H. pylori* strains used in this study are listed in **Supplemental Table 1**. General culture of all strains was performed on solid media using Columbia agar plates containing 4% Columbia agar base (Oxoid), 5% defibrinated horse blood (Hemostat Labs), 10 mg/mL vancomycin (Thermo Fisher), 2.5 U/mL polymyxin B (Sigma-Aldrich), 8 mg/mL amphotericin B (Sigma-Aldrich), and 0.2% β-cyclodextrin (Thermo Fisher). Plates used to grow *H. pylori* from homogenized mouse stomach also included 5 mg/L cefsulodin (Thermo Fisher), 5 mg/L trimethoprim (Sigma) and 0.2 mg/mL of bacitracin (Acros Organics, Fisher) to prevent outgrowth of mouse commensal bacteria and fungi. For murine competition experiments, 25 μg/mL kanamycin was added to distinguish between cassette-marked (kanamycin-resistant) and unmarked (kanamycin-sensitive) bacteria. Liquid cultures, when required, were grown shaking in Brucella broth (BD BBL) supplemented with 10% heat-inactivated FBS (Innovative Bioscience). *H. pylori* on plates and in liquid culture were grown at 37 °C in microaerobic conditions (10% CO_2_, 10% O_2_, and 80% N_2_).

### Biotinylation of *H. pylori* for BASEHIT profiling

*H. pylori* grown in liquid culture was biotinylated as previously described (**39**). Briefly, five optical density at 600 nm (OD_600_) units of culture were collected by centrifugation at 1,500 *g* (Eppendorf 5424) for five minutes, then washed in PBS (Gibco) three times. The pellet was then resuspended in 5 uM EZ-Link™ Sulfo-NHS-Biotin (Thermo Scientific) in PBS and incubated at 37 ⁰C for thirty minutes. The reaction was stopped with 5 µL of 1M Tris-HCl, pH 8.0. The cells were then washed once with PBS and frozen in PBS with 10% glycerol at a final concentration of five OD_600_/mL until use.

### BASEHIT yeast library selection

BASEHIT profiling was performed as previously described (**39**). The yeast library was cultured in SDO-Ura at 30 °C then, induced by resuspension at an OD_600_ of 1 in SGO-Ura supplemented with 10% SDO-Ura for 24 hours. Plasmid DNA was extracted from 400 µl of the pre-selection library, to allow for comparison to post-selection libraries, using a Zymoprep Yeast Plasmid Miniprep II kit (Zymo Research). Induced yeast cells (3·10^7^) were pelleted in 96-well v-bottom microtitre plates, then resuspended in 100 µl PBE (PBS, 0.5% BSA, 0.5 mM EDTA). Then 50 µl of five OD_600_/mL biotinylated bacteria were added, and the yeast were incubated with shaking for 2 hours at 4 °C. Yeast were washed once with 200 µl PBE, resuspended in 100 µl PBE with a 1:100 dilution of streptavidin microparticles (0.29 µm; Spherotech) and incubated with shaking for 1 hour at 4 °C. Yeast were washed once with 200 µl PBE, then pelleted and resuspended in 150 µl PBE at room temperature. A 96-well magnet was used to remove bead-bound yeast, which were washed twice with 150 µl PBE. Washed yeast cells were eluted into 150 µl PBE by removal of the magnet. Selected yeast were pelleted and expanded by growth in 1 ml SDO-Ura supplemented with chloramphenicol at 30 °C for 44 hours, before having DNA extracted for sequencing.

### Yeast barcode sequencing

BASEHIT yeast library barcodes were sequenced as previously described (**39**). DNA was extracted from yeast libraries using Zymoprep Yeast Plasmid Miniprep II kits (Zymo Research). The protein display barcode on the yeast plasmid was amplified through PCR with the primers 159_DIF2 and 159_DIR2. The PCR product was directly used as template for the second round of PCR using Nextera i5 and i7 dual-index library primers (Illumina). PCR products were pooled and run on a 2% agarose gel, then purified using the QIAquick Gel Extraction Kit (Qiagen). The next-generation sequencing library was sequenced by the Yale Center for Genome Analysis (YCGA), using a full Illumina NovaSeq S4 2 × 150 lane to obtain 2 billion total reads.

### Computational analysis of BASEHIT results

The raw BASEHIT barcode sequences were analyzed by the basehitmodel R pipeline (**39**) to generate interaction (ixn) scores and binding hit calls. Samples were demultiplexed and sequenced barcodes were mapped, only accepting exact barcode matches to ensure that correct calls were used in the analysis. The input to the statistical model consists of counts of barcodes distributed across 3,324 human proteins in an input sample, three beads-only samples and three output samples for each strain. The basehitmodel function model_proteins_separately was used with arguments iter_sampling = 2500, iter_warmup = 2000, algorithm = ‘mcmc’, seed = 123, run in parallel across eight cores. This model uses a zero-inflated negative binomial model to convert barcode reads into the posterior mean calculated for each protein separately, accounting for differences in the variance in input and general nonspecific binding in each. After the model for each protein is established, the concordance over the three replicates is calculated for each sample as a measure of consistency of effect. The model derives the interaction score from the posterior mean, then uses the calculated posterior interval, magnitude of the score, and concordance to establish binding hits based on standard (95% interval excludes zero, estimated effect size > 0.5, and concordance score > 0.75) or stringent (95% interval excludes zero, estimated effect size > 0.5, and concordance score > 0.95) thresholds favoring large, consistent effect and inter-replicate concordances.

The ixn score data were then used to produce hierarchical clustering using Seaborn, principal component analysis with matplotlib (https://github.com/matplotlib/matplotlib) and sklearn (https://github.com/scikit-learn/scikit-learn), and ANOVA with scipy (https://github.com/scipy/scipy). Normalized ixn scores per strain were generated by subtracting the lowest ixn score per strain from each score in that strain, then dividing them by the range for that strain. For the correlation analysis, a Pearson correlation matrix of unnormalized ixn scores was generated in R and aligned to an ANI matrix generated from Anvi’o (https://github.com/merenlab/anvio) (**72**) using the vegan package (https://vegandevs.github.io/vegan/) to generate aligned vectors and R to calculate the overall Pearson correlation between the two matrices.

### Genome wide association studies

Genome wide association studies were performed using bugwas (https://github.com/sgearle/bugwas) and pyseer (https://github.com/mgalardini/pyseer) on the ixn scores for each human exoprotein against SNPs and InDels called for J99 culture collection whole genome sequence data (BioProject: PRJNA622860), as determined by Breseq (https://github.com/barricklab/breseq) (for bugwas) or Snippy (https://github.com/tseemann/snippy) (for pyseer) against the reference J99 sequence (GenBank: AE001439.1). Results were then filtered to remove p-values above 0.05 (insignificant) and p-values that were repeated more than five times within a single exoprotein’s GWAS (likely present due to lineage). The results were then combined into a single list of loci and their relationship to ixn score differences for specific exoproteins. A more stringent p-value cutoff of - log(p-value) = 4 (derived from 1/number of polymorphisms tested) was applied for visualization and inclusion in **Table 1**. These results were then annotated with BioPython (https://github.com/biopython/biopython) and graphed as a Manhattan plot with matplotlib.

### Adherence of fluorescein-labeled *H. pylori* to *ex vivo* tissue

The *ex vivo* tissue adherence assay was performed as previously described (**73**). Briefly, five OD_600_ units of mid-log liquid *H. pylori* culture are pelleted by centrifugation at 1400 *g* (Tx-160 centrifuge with TMY-27 rotor) for five minutes, then washed in PBS-T (PBS with 0.05% Tween-20). Bacteria were washed once in carbonate buffer (100 mM sodium carbonate, 150 mM NaCl, pH 9.0), then resuspended in carbonate buffer with 10 µg/mL fluorescein isothiocyanate (FITC, Sigma) and incubated rotating for ten minutes, protected from light. Bacteria were then collected by centrifugation at 1500 *g* for five minutes and washed in blocking buffer (1% periodate-oxidized bovine serum albumin in PBS-T) three times. The concentration was adjusted to one OD_600_/mL in blocking buffer, aliquoted, and then stored at -70 ⁰C until use. Formalin fixed paraffin embedded (FFPE) mouse gastric tissue slices (5 µm thick) were prepared by deparaffinization and rehydration using Histo-Clear (National Diagnostics) twice for ten minutes, then once for three minutes, followed by three five-minute washes in 100% isopropanol, one three-minute wash each in 95% ethanol then 50% ethanol, five minutes in deionized water, and finally three five-minute washes in PBS. Stomachs were circled with a wax pen, then blocked for 2.5 hours with blocking buffer. After removing the blocking buffer, FITC-labeled *H. pylori* were added at OD_600_ = 0.05 in blocking buffer. The bacteria were allowed to adhere for two hours in a hydration chamber, protected from light. The slides were washed with three rounds of dunking, 30 times each in PBS-T, then mounted with ProLong Diamond antifade mountant with DAPI (Life Technologies). Slides were then visualized by fluorescent slide scanning microscopy on TissueFAXS (TissueGnostics) or Evident VS200 (Olympus Evident) then analyzed on Imaris (Oxford Instruments) to calculate the fluorescent spots (proxy for bacteria) per area (µm^2^) of DAPI-positive glandular stomach tissue.

### Fast Lipid Analysis Technique (FLAT)

FLAT was performed as previously described (**51**). Briefly, 1 μL of the prepared citrate buffer solution (0.2 M citric acid, 0.1 M trisodium citrate, pH 3.8) was deposited onto the sample spot on the 96-well MALDI plate. The plate was incubated in a humidified, closed heat block (panini press) for 30 min at 110° C. After heating, the MALDI plate was removed from the panini press, cooled, and the plate was thoroughly washed with water using a pipettor several times and left to dry on the laboratory bench. Bruker microflex and (timsTOF) flex have been employed for rapid profiling lipid A and structural analysis by tandem mass spectrometry (FLAT) (**74**). The Bruker microflex (LR) was conducted in reflectron-negative ion modes. Spectra were collected using 50% global intensity with approximately 300 laser shots per spectrum. All spectra were collected between *m/z* 1,000 and 2,400. Mass calibration was performed using an electrospray tuning mix (Agilent, Palo Alto, CA). Bruker (timsTOF) MS was equipped with a dual ESI/MALDI source with a SmartBeam 3D 10 KHz frequency tripled Nd:YAG laser (355 nm). The system was operated in “qTOF” mode (TIMS deactivated). Ion transfer tuning was used with the following parameters: Funnel 1 RF: 440.0 Vpp, Funnel 2 RF: 490.0 Vpp, Multipole RF 490.0 Vpp, is CID Energy: 0.0 eV, and Deflection Delta: -60.0 V. Quadrupole has been used with the following values for MS mode: Ion Energy: 4.0 eV and low mass 700.00 *m/z*. Collision cell activation of ions used the following values for MS mode. Collision Energy: 9.0 eV and Collision RF: 3900.0 Vpp. In the MS/MS mode, the precursor ion was chosen by typing the targeted *m/z* value, including two digits to the right of the decimal point. Typical isolation width and collision energy were set to 4 - 6 *m/z* and 100 - 110 eV, respectively. Focus Pre TOF used the following values for Transfer Time 110.0 µs and Pre-Pulse Storage 9.0 µs. Agilent ESI Tune Mix was used to perform calibration of the *m/z* scale. MALDI parameters in qTOF were optimized to maximize intensity by tuning ion optics, laser intensity, and laser focus. All mass spectra were collected at 104 µm laser diameter with beam scan on using 800 laser shots per spot and 70 - 80% laser power. Both MS and MS/MS data were collected in negative ion mode. All MALDI (timsTOF) MS and MS/MS data were visualized using mMass (Ver 5.5.0) (**75**). Peak picking was conducted in mMass. Identification of all fragment ions was determined based on Chemdraw Ultra (Ver23.1.1).

### LPS extraction and qualitative analysis with SDS-PAGE

LPS extraction was performed by warm phenol/ether extraction as described by Davis and Goldberg (**76**). Briefly, 5-10 OD_600_ units of *H. pylori* were collected from mid-log phase by centrifugation. The pellet was then resuspended in 200 µL of 1x SDS buffer (2% β-mercaptoethanol, 2% sodium dodecyl sulfate, a pinch of bromophenol blue, and 10% glycerol in 0.05 M Tris-HCl, pH 6.8). The bacteria were boiled at 100 ⁰C for 15 minutes and then cooled to room temperature. Proteins were degraded by adding 10 µL of 10 mg/mL proteinase K and incubation at 59 ⁰C overnight. Next, 200 µL of ice-cold Tris-saturated phenol was added and vortexed for ten seconds, then incubated shaking vigorously for fifteen minutes at 65 ⁰C. After cooling to room temperature, 1 mL of diethyl ether was added and then vortexed for five seconds. Samples were spun at 20,600 *g* (Eppendorf 5424) for ten minutes to separate phases. The bottom blue layer was saved, then re-extracted with an additional round of phenol/ether as above. The resulting samples were then added to 200 µL of a 2x SDS buffer. Samples were then run on 4-15% gradient SDS-PAGE Tris-glycine gels (Bio Rad Mini-PROTEAN TGX) with discontinuous running buffer (cathode: 0.1 M Tris, 0.1 M glycine, 0.1% SDS, pH 8.25; anode: 0.2 M Tris-HCl, pH 8.9) at 100 V until the dye front was near the bottom. Gels were then fixed overnight in 60% methanol and 10% glacial acetic acid, then washed in 3% acetic acid for twenty minutes with rocking. The carbohydrates were then oxidized with 1% periodic acid in 3% acetic acid for thirty minutes. Gels were washed three times in 3% acetic acid for twenty minutes each, then treated with ProQ Emerald 300 Lipopolysaccharide kit (Molecular Probes) stain in staining solution for two hours protected from light. Gels were washed two times with 3% acetic acid for twenty minutes, then visualized using a Bio Rad ChemiDoc system.

### Whole genome sequencing with Oxford Nanopore

High quality *H. pylori* genomic DNA was extracted with a CsCl gradient to prevent shearing. Two full plates of healthy mid-log bacteria were scraped and resuspended in 3.5 mL of STE (0.1 M NaCl, 10 mM Tris-HCl, 1 mM EDTA, pH 8.0) with lysozyme (350 µg/mL), then incubated at 37 ⁰C for fifteen minutes. Complete lysis was then performed by adding 350 µL of 10% SDS and incubating at 65 ⁰C for 15 minutes. Proteins were digested with 35 µL of proteinase K (50 mg/mL) at 50 ⁰C for 2.5 hours. Solid CsCl was added to a final concentration of 1 g/mL, mixed by inversion, then incubated at 65 ⁰C for fifteen minutes. Ethidium bromide (10 mg/mL) was added at 80 µg/mL, then incubated at 65 ⁰C for forty minutes. Samples were transferred to g-Max Quick-Seal tubes (Beckman Coulter) and centrifuged at 390,000 *g* (Beckman Coulter - Type 70.1Ti rotor) at 20 ⁰C overnight. DNA was visualized with UV and carefully extracted from the tubes with a 20-gauge needle and syringe. DNA was then cleaned with butanol extraction five times with one volume of STE-saturated butanol. DNA was then precipitated with five volumes of 100% ethanol. DNA strands were moved to TE (10 mM Tris-HCl, 100 mM EDTA, pH 8.0) for storage using flame-closed glass Pasteur pipettes and allowed to rehydrate at 4 ⁰C overnight before freezing. Samples were quantified and quality checked by Tapestation (Agilent 4200), then submitted for standard bacterial genome sequencing through Plasmidsaurus.

### Genetic analysis

Analysis of previously sequenced strains was performed by retrieval of the sequences from the NCBI database and Mauve alignment in Geneious. Analysis of newly generated Oxford Nanopore sequencing was performed by using reference guided assembly (to G27 (GenBank: CP001173.1) for NSH57 or PMSS1 (GenBank: CP018823.1) for the new mouse output isolates) through wf-bacterial-genomes in Epi2me to generate both variant calls and draft genomes for Mauve alignment in Geneious.

### Mouse coinfection by oral gavage

Mice were infected and CFU elucidated as previously described (**20**). Ten female C57BL/6NJ mice (Jackson Laboratories, stock number 005304) per group were inoculated by oral gavage with 5·10^7^ CFU total of an equal mixture of PMSS1 and one-day PMSS1 mouse infection output isolates (G, H, and I in Fig. 4C) which had been transformed with a kanamycin cassette (*aphA3*) containing plasmid pDCY40 (**77**) to insert the kanamycin resistance gene at a neutral locus between *HpG27_0186 (HPYLPMSS1_0192)* and *HpG27_0187 (HPYLPMSS1_0193)*. Likewise, 5 male C57BL/6NJ mice were inoculated with the same mixtures, but the cassette was now moved to a similarly constructed parental PMSS1 strain and not in G, H, or I. After one week of colonization, mice were euthanized. Stomachs were excised and forestomaches removed. They were then opened along the lesser curvature and food contents were gently scraped. Half of the processed stomach, containing both antrum and corpus, was weighed and placed in 500 µL of liquid *H. pylori* media for mechanical homogenization with pellet pestles (DWK Life Sciences). Serial dilutions were then plated on both plain and kanamycin-containing media. Plates were incubated for 5 to 7 days, then the CFU were enumerated.

### DNA extraction from mouse gastric homogenates and amplicon sequencing

DNA was extracted from mouse gastric homogenates by phenol/chloroform extraction (**78**). Briefly, 700 µL lysis buffer (10 mM Tris-HCl, 10 mM EDTA, 15 mg/mL lysozyme, and 100 µg/mL RNase A, pH 8.0) was added to 300 µL of homogenized mouse stomach halves suspended in liquid *H. pylori* media, then incubated shaking at 37 ⁰C for one hour. Next, 30 µL of 10% SDS and 3 µL of 20 mg/mL proteinase K were added, then incubated at 55 ⁰C for one hour. Then, 525 µL of phenol:chloroform:isoamyl alcohol (25:24:1) was added and mixed by inversion for ten minutes at room temperature. Phase separation was established by centrifugation at 13,500 *g* (Eppendorf 5424) for fifteen minutes. The upper aqueous phase was then added to 1 mL of -20 ⁰C 100% ethanol. DNA was pelleted by centrifugation at 13,500 *g* for 15 minutes. DNA pellets were air dried, then rehydrated in TE (10 mM Tris-HCl, 100 mM EDTA, pH 8.0). The *futB* locus was then amplified with primers JPF279 (5’-AAATGCGACTCTAACAGCCTTAATTTAGAAG-3’) and JPF282 (5’-GATCGTGGAAAATGACGCCTTAAACGGC-3’). Purified PCRs were then sequenced using the Plasmidsaurus premium PCR pipeline. Sequencing data was then analyzed with the Epi2me wf-amplicon pipeline and Geneious.

## Data Availability

Sequencing data produced in this study has been deposited in the NCBI Short Read Archive (BioProject: PRJNA1430809).

## Acknowledgements

This research was supported by NIH R01 AI054423 (NRS), RO1 AI147314 (RKE), R01 AI104895 (RKE), and The Leona M. and Harry B. Helmsley Charitable Trust (NWP) and used resources from the NIH P30 CA015704 of the Fred Hutch/University of Washington/Seattle Children’s Cancer Consortium, which includes the Cellular Imaging, Comparative Medicine, and Experimental Histopathology Shared Resources.

## Author Contributions

Based on CRediT Taxonomy

**JPF**: Conceptualization, Validation, Formal Analysis, Investigation, Data Curation, Writing – Original Draft, Visualization, Project administration; **NDS:** Methodology, Formal Analysis, Investigation, Writing - Review & Editing; **JAS:** Investigation, Writing - Review & Editing; **MO:** Investigation, Formal Analysis, Writing - Review & Editing, Visualization; **HY:** Investigation, Formal Analysis, Writing - Review & Editing, Visualization; **MNS:** Investigation, Writing - Review & Editing; **AMR:** Conceptualization, Methodology, Resources, Writing - Review & Editing, Supervision, Funding acquisition; **RKE:** Conceptualization, Methodology, Resources, Writing - Review & Editing, Supervision, Funding acquisition; **NWP:** Conceptualization, Methodology, Resources, Writing - Review & Editing, Supervision, Funding acquisition; **NRS:** Conceptualization, Methodology, Resources, Data Curation, Writing – Original Draft, Supervision, Project administration, Funding acquisition

## References

1. Chen YC, Malfertheiner P, Yu HT, Kuo CL, Chang YY, Meng FT, Wu YX, Hsiao JL, Chen MJ, Lin KP, Wu CY, Lin JT, O’Morain C, Megraud F, Lee WC, El-Omar EM, Wu MS, Liou JM. 2024. Global Prevalence of Helicobacter pylori Infection and Incidence of Gastric Cancer Between 1980 and 2022. Gastroenterology. 166:605–19.

2. Plummer M, Franceschi S, Vignat J, Forman D, de Martel C. 2015. Global burden of gastric cancer attributable to Helicobacter pylori. Int J Cancer. 136:487–90.

3. de Martel C, Georges D, Bray F, Ferlay J, Clifford GM. 2020. Global burden of cancer attributable to infections in 2018: a worldwide incidence analysis. Lancet Glob Health. 8:e180–e90.

4. Alm RA, Ling LS, Moir DT, King BL, Brown ED, Doig PC, Smith DR, Noonan B, Guild BC, deJonge BL, Carmel G, Tummino PJ, Caruso A, Uria-Nickelsen M, Mills DM, Ives C, Gibson R, Merberg D, Mills SD, Jiang Q, Taylor DE, Vovis GF, Trust TJ. 1999. Genomic-sequence comparison of two unrelated isolates of the human gastric pathogen Helicobacter pylori. Nature. 397:176–80.

5. Alm RA, Trust TJ. 1999. Analysis of the genetic diversity of Helicobacter pylori: the tale of two genomes. J Mol Med. 77:834–46.

6. Draper JL, Hansen LM, Bernick DL, Abedrabbo S, Underwood JG, Kong N, Huang BC, Weis AM, Weimer BC, van Vliet AH, Pourmand N, Solnick JV, Karplus K, Ottemann KM. 2017. Fallacy of the Unique Genome: Sequence Diversity within Single Helicobacter pylori Strains. mBio. 8:

7. Gressmann H, Linz B, Ghai R, Pleissner KP, Schlapbach R, Yamaoka Y, Kraft C, Suerbaum S, Meyer TF, Achtman M. 2005. Gain and loss of multiple genes during the evolution of Helicobacter pylori. PLoS Genet. 1:e43.

8. Jackson LK, Potter B, Schneider S, Fitzgibbon M, Blair K, Farah H, Krishna U, Bedford T, Peek Jr RM, Salama NR. 2020. Helicobacter pylori diversification during chronic infection within a single host generates sub-populations with distinct phenotypes. PLoS pathogens. 16:e1008686.

9. Suerbaum S, Josenhans C. 2007. Helicobacter pylori evolution and phenotypic diversification in a changing host. Nat Rev Microbiol. 5:441–52.

10. Israel DA, Salama N, Krishna U, Rieger UM, Atherton JC, Falkow S, Peek RM, Jr. 2001. Helicobacter pylori genetic diversity within the gastric niche of a single human host. Proc Natl Acad Sci U S A. 98:14625–30.

11. Alm RA, Bina J, Andrews BM, Doig P, Hancock RE, Trust TJ. 2000. Comparative genomics of Helicobacter pylori: analysis of the outer membrane protein families. Infect Immun. 68:4155–68.

12. Colbeck JC, Hansen LM, Fong JM, Solnick JV. 2006. Genotypic profile of the outer membrane proteins BabA and BabB in clinical isolates of Helicobacter pylori. Infect Immun. 74:4375–8.

13. O’Brien VP, Jackson LK, Frick JP, Martinez AER, Jones DS, Johnston CD, Salama NR. 2023. Helicobacter pylori Chronic Infection Selects for Effective Colonizers of Metaplastic Glands. mBio. 14:e03116–22.

14. Backstrom A, Lundberg C, Kersulyte D, Berg DE, Boren T, Arnqvist A. 2004. Metastability of Helicobacter pylori bab adhesin genes and dynamics in Lewis b antigen binding. Proc Natl Acad Sci U S A. 101:16923–8.

15. Doig P, Trust TJ. 1994. Identification of surface-exposed outer membrane antigens of Helicobacter pylori. Infect Immun. 62:4526–33.

16. Ilver D, Arnqvist A, Ogren J, Frick IM, Kersulyte D, Incecik ET, Berg DE, Covacci A, Engstrand L, Boren T. 1998. Helicobacter pylori adhesin binding fucosylated histo-blood group antigens revealed by retagging. Science. 279:373–7.

17. Boren T, Falk P, Roth KA, Larson G, Normark S. 1993. Attachment of Helicobacter pylori to human gastric epithelium mediated by blood group antigens. Science (Washington D C). 262:1892–95.

18. Mahdavi J, Sonden B, Hurtig M, Olfat FO, Forsberg L, Roche N, Angstrom J, Larsson T, Teneberg S, Karlsson KA, Altraja S, Wadstrom T, Kersulyte D, Berg DE, Dubois A, Petersson C, Magnusson KE, Norberg T, Lindh F, Lundskog BB, Arnqvist A, Hammarstrom L, Boren T. 2002. Helicobacter pylori SabA adhesin in persistent infection and chronic inflammation. Science. 297:573–8.

19. Javaheri A, Kruse T, Moonens K, Mejias-Luque R, Debraekeleer A, Asche CI, Tegtmeyer N, Kalali B, Bach NC, Sieber SA, Hill DJ, Koniger V, Hauck CR, Moskalenko R, Haas R, Busch DH, Klaile E, Slevogt H, Schmidt A, Backert S, Remaut H, Singer BB, Gerhard M. 2016. Helicobacter pylori adhesin HopQ engages in a virulence-enhancing interaction with human CEACAMs. Nat Microbiol. 2:16189.

20. O’Brien VP, Koehne AL, Dubrulle J, Rodriguez AE, Leverich CK, Kong VP, Campbell JS, Pierce RH, Goldenring JR, Choi E, Salama NR. 2021. Sustained Helicobacter pylori infection accelerates gastric dysplasia in a mouse model. Life Sci Alliance. 4:

21. Talarico S, Whitefield SE, Fero J, Haas R, Salama NR. 2012. Regulation of Helicobacter pylori adherence by gene conversion. Mol Microbiol.

22. Aspholm-Hurtig M, Dailide G, Lahmann M, Kalia A, Ilver D, Roche N, Vikstrom S, Sjostrom R, Linden S, Backstrom A, Lundberg C, Arnqvist A, Mahdavi J, Nilsson UJ, Velapatino B, Gilman RH, Gerhard M, Alarcon T, Lopez-Brea M, Nakazawa T, Fox JG, Correa P, Dominguez-Bello MG, Perez-Perez GI, Blaser MJ, Normark S, Carlstedt I, Oscarson S, Teneberg S, Berg DE, Boren T. 2004. Functional adaptation of BabA, the H. pylori ABO blood group antigen binding adhesin. Science. 305:519–22.

23. Hanisch FG, Bonar D, Schloerer N, Schroten H. 2014. Human trefoil factor 2 is a lectin that binds α-GlcNAc-capped mucin glycans with antibiotic activity against Helicobacter pylori. J Biol Chem. 289:27363–75.

24. Clyne M, Drumm B. 1997. Absence of effect of Lewis a and Lewis b expression on adherence of Helicobacter pylori to human gastric cells. Gastroenterology. 113:72–80.

25. Clyne M, Drumm B. 1996. Cell envelope characteristics of Helicobacter pylori: their role in adherence to mucosal surfaces and virulence. FEMS Immunol Med Microbiol. 16:141–55.

26. Reeves EP, Ali T, Leonard P, Hearty S, O’Kennedy R, May FE, Westley BR, Josenhans C, Rust M, Suerbaum S, Smith A, Drumm B, Clyne M. 2008. Helicobacter pylori lipopolysaccharide interacts with TFF1 in a pH-dependent manner. Gastroenterology. 135:2043–54, 54.e1-2.

27. Jacquies M. 1996. Role of lipo-oligosaccharides and lipopolysaccharides in bacterial adherence. Trends in Microbiology. 4:408–10.

28. Moran AP. 1996. The role of lipopolysaccharide in Helicobacter pylori pathogenesis. Aliment Pharmacol Ther. 10 Suppl 1:39–50.

29. Mahdavi J, Boren T, Vandenbroucke-Grauls C, Appelmelk BJ. 2003. Limited role of lipopolysaccharide Lewis antigens in adherence of Helicobacter pylori to the human gastric epithelium. Infect Immun. 71:2876–80.

30. Tang X, Yang T, Shen Y, Song X, Benghezal M, Marshall BJ, Tang H, Li H. 2023. Roles of Lipopolysaccharide Glycosyltransferases in Maintenance of Helicobacter pylori Morphology, Cell Wall Permeability, and Antimicrobial Susceptibilities. Int J Mol Sci. 24:

31. Li H, Tang X, Yang T, Liao T, Debowski AW, Yang T, Shen Y, Nilsson HO, Haslam SM, Mulloy B, Dell A, Stubbs KA, Fischer W, Haas R, Tang H, Marshall BJ, Benghezal M. 2025. Reinvestigation into the role of lipopolysaccharide Glycosyltransferases in Helicobacter pylori protein glycosylation. Gut Microbes. 17:2455513.

32. Li H, Marceau M, Yang T, Liao T, Tang X, Hu R, Xie Y, Tang H, Tay A, Shi Y, Shen Y, Yang T, Pi X, Lamichhane B, Luo Y, Debowski AW, Nilsson HO, Haslam SM, Mulloy B, Dell A, Stubbs KA, Marshall BJ, Benghezal M. 2019. East-Asian Helicobacter pylori strains synthesize heptan-deficient lipopolysaccharide. PLoS Genet. 15:e1008497.

33. Appelmelk BJ, Martin SL, Monteiro MA, Clayton CA, McColm AA, Zheng P, Verboom T, Maaskant JJ, van den Eijnden DH, Hokke CH, Perry MB, Vandenbroucke-Grauls CM, Kusters JG. 1999. Phase variation in Helicobacter pylori lipopolysaccharide due to changes in the lengths of poly(C) tracts in alpha3-fucosyltransferase genes. Infect Immun. 67:5361–6.

34. Salaun L, Linz B, Suerbaum S, Saunders NJ. 2004. The diversity within an expanded and redefined repertoire of phase-variable genes in Helicobacter pylori. Microbiology. 150:817–30.

35. Salaün L, Saunders NJ. 2006. Population-associated differences between the phase variable LPS biosynthetic genes of Helicobacter pylori. BMC Microbiol. 6:79.

36. Wang G, Rasko DA, Sherburne R, Taylor DE. 1999. Molecular genetic basis for the variable expression of Lewis Y antigen in Helicobacter pylori: analysis of the alpha (1,2) fucosyltransferase gene. Mol Microbiol. 31:1265–74.

37. Nilsson C, Skoglund A, Moran AP, Annuk H, Engstrand L, Normark S. 2006. An enzymatic ruler modulates Lewis antigen glycosylation of Helicobacter pylori LPS during persistent infection. Proc Natl Acad Sci U S A. 103:2863–8.

38. Nilsson C, Skoglund A, Moran AP, Annuk H, Engstrand L, Normark S. 2008. Lipopolysaccharide diversity evolving in Helicobacter pylori communities through genetic modifications in fucosyltransferases. PLoS One. 3:e3811.

39. Sonnert ND, Rosen CE, Ghazi AR, Franzosa EA, Duncan-Lowey B, González-Hernández JA, Huck JD, Yang Y, Dai Y, Rice TA, Nguyen MT, Song D, Cao Y, Martin AL, Bielecka AA, Fischer S, Guan C, Oh J, Huttenhower C, Ring AM, Palm NW. 2024. A host-microbiota interactome reveals extensive transkingdom connectivity. Nature. 628:171–79.

40. Lee A, O’Rourke J, De Ungria MC, Robertson B, Daskalopoulos G, Dixon MF. 1997. A standardized mouse model of Helicobacter pylori infection: Introducing the Sydney strain. Gastroenterology. 112:1386–97.

41. Covacci A, Censini S, Bugnoli M, Petracca R, Burroni D, Macchia G, Massone A, Papini E, Xiang Z, Figura N, et al. 1993. Molecular characterization of the 128-kDa immunodominant antigen of Helicobacter pylori associated with cytotoxicity and duodenal ulcer. Proc Natl Acad Sci U S A. 90:5791–5.

42. Tomb JF, White O, Kerlavage AR, Clayton RA, Sutton GG, Fleischmann RD, Ketchum KA, Klenk HP, Gill S, Dougherty BA, Nelson K, Quackenbush J, Zhou L, Kirkness EF, Peterson S, Loftus B, Richardson D, Dodson R, Khalak HG, Glodek A, McKenney K, Fitzegerald LM, Lee N, Adams MD, Venter JC, et al. 1997. The complete genome sequence of the gastric pathogen Helicobacter pylori. Nature. 388:539–47.

43. Baldwin DN, Shepherd B, Kraemer P, Hall MK, Sycuro LK, Pinto-Santini DM, Salama NR. 2007. Identification of Helicobacter pylori genes that contribute to stomach colonization. Infect Immun. 75:1005–16.

44. Dunne C, Naughton J, Duggan G, Loughrey C, Kilcoyne M, Joshi L, Carrington S, Earley H, Backert S, Robbe Masselot C, May FEB, Clyne M. 2018. Binding of Helicobacter pylori to Human Gastric Mucins Correlates with Binding of TFF1. Microorganisms. 6:

45. Yang DC, Blair KM, Taylor JA, Petersen TW, Sessler T, Tull CM, Leverich CK, Collar AL, Wyckoff TJ, Biboy J, Vollmer W, Salama NR. 2019. A Genome-Wide Helicobacter pylori Morphology Screen Uncovers a Membrane-Spanning Helical Cell Shape Complex. J Bacteriol. 201:

46. Earle SG, Wu CH, Charlesworth J, Stoesser N, Gordon NC, Walker TM, Spencer CCA, Iqbal Z, Clifton DA, Hopkins KL, Woodford N, Smith EG, Ismail N, Llewelyn MJ, Peto TE, Crook DW, McVean G, Walker AS, Wilson DJ. 2016. Identifying lineage effects when controlling for population structure improves power in bacterial association studies. Nat Microbiol. 1:16041.

47. Lees JA, Galardini M, Bentley SD, Weiser JN, Corander J. 2018. pyseer: a comprehensive tool for microbial pangenome-wide association studies. Bioinformatics. 34:4310–12.

48. Guo S, Skala W, Magdolen V, Brandstetter H, Goettig P. 2014. Sweetened kallikrein-related peptidases (KLKs): glycan trees as potential regulators of activation and activity. Biol Chem. 395:959–76.

49. Kaskow BJ, Proffitt JM, Blangero J, Moses EK, Abraham LJ. 2012. Diverse biological activities of the vascular non-inflammatory molecules - the Vanin pantetheinases. Biochem Biophys Res Commun. 417:653–8.

50. Mori Y, Hamuro T, Nakashima T, Hamamoto T, Natsuka S, Hase S, Iwanaga S. 2009. Biochemical characterization of plasma-derived tissue factor pathway inhibitor: post-translational modification of free, full-length form with particular reference to the sugar chain. J Thromb Haemost. 7:111–20.

51. Sorensen M, Chandler CE, Gardner FM, Ramadan S, Khot PD, Leung LM, Farrance CE, Goodlett DR, Ernst RK, Nilsson E. 2020. Rapid microbial identification and colistin resistance detection via MALDI-TOF MS using a novel on-target extraction of membrane lipids. Sci Rep. 10:21536.

52. Li H, Yang T, Liao T, Debowski AW, Nilsson HO, Fulurija A, Haslam SM, Mulloy B, Dell A, Stubbs KA, Marshall BJ, Benghezal M. 2017. The redefinition of Helicobacter pylori lipopolysaccharide O-antigen and core-oligosaccharide domains. PLoS Pathog. 13:e1006280.

53. Saunders NJ, Peden JF, Hood DW, Moxon ER. 1998. Simple sequence repeats in the Helicobacter pylori genome. Mol Microbiol. 27:1091–8.

54. Choi E, Means AL, Coffey RJ, Goldenring JR. 2019. Active Kras Expression in Gastric Isthmal Progenitor Cells Induces Foveolar Hyperplasia but Not Metaplasia. Cell Mol Gastroenterol Hepatol. 7:251–53 e1.

55. Moran AP, Lindner B, Walsh EJ. 1997. Structural characterization of the lipid A component of Helicobacter pylori rough- and smooth-form lipopolysaccharides. J Bacteriol. 179:6453–63.

56. Suda Y, Ogawa T, Kashihara W, Oikawa M, Shimoyama T, Hayashi T, Tamura T, Kusumoto S. 1997. Chemical structure of lipid A from Helicobacter pylori strain 206-1 lipopolysaccharide. J Biochem. 121:1129–33.

57. Suda Y, Kim YM, Ogawa T, Yasui N, Hasegawa Y, Kashihara W, Shimoyama T, Aoyama K, Nagata K, Tamura T, Kusumoto S. 2001. Chemical structure and biological activity of a lipid A component from Helicobacter pylori strain 206. J Endotoxin Res. 7:95–104.

58. Tang X, Tay A, Benghezal M, Marshall BJ, Tang H, Li H. 2025. Advances in Helicobacter pylori lipopolysaccharide structure and function. FEMS Microbiol Rev. 49:

59. Altman E, Chandan V, Harrison BA, Vinogradov E. 2018. Structural and immunological characterization of a glycoconjugate based on the delipidated lipopolysaccharide from a nontypeable Helicobacter pylori strain PJ1 containing an extended d-glycero-d-manno-heptan. Carbohydr Res. 456:19–23.

60. Tran AX, Karbarz MJ, Wang X, Raetz CR, McGrath SC, Cotter RJ, Trent MS. 2004. Periplasmic cleavage and modification of the 1-phosphate group of Helicobacter pylori lipid A. J Biol Chem. 279:55780–91.

61. Tran AX, Whittimore JD, Wyrick PB, McGrath SC, Cotter RJ, Trent MS. 2006. The lipid A 1-phosphatase of Helicobacter pylori is required for resistance to the antimicrobial peptide polymyxin. J Bacteriol. 188:4531–41.

62. Cullen TW, Giles DK, Wolf LN, Ecobichon C, Boneca IG, Trent MS. 2011. Helicobacter pylori versus the host: remodeling of the bacterial outer membrane is required for survival in the gastric mucosa. PLoS Pathog. 7:e1002454.

63. Salaun L, Ayraud S, Saunders NJ. 2005. Phase variation mediated niche adaptation during prolonged experimental murine infection with Helicobacter pylori. Microbiology. 151:917–23.

64. Solnick JV, Hansen LM, Salama NR, Boonjakuakul JK, Syvanen M. 2004. Modification of Helicobacter pylori outer membrane protein expression during experimental infection of rhesus macaques. Proc Natl Acad Sci U S A. 101:2106–11.

65. Aras RA, Fischer W, Perez-Perez GI, Crosatti M, Ando T, Haas R, Blaser MJ. 2003. Plasticity of repetitive DNA sequences within a bacterial (Type IV) secretion system component. J Exp Med. 198:1349–60.

66. Barrozo RM, Hansen LM, Lam AM, Skoog EC, Martin ME, Cai LP, Lin Y, Latoscha A, Suerbaum S, Canfield DR, Solnick JV. 2016. CagY Is an Immune-Sensitive Regulator of the Helicobacter pylori Type IV Secretion System. Gastroenterology. 151:1164–75 e3.

67. Barrozo RM, Cooke CL, Hansen LM, Lam AM, Gaddy JA, Johnson EM, Cariaga TA, Suarez G, Peek RM, Jr., Cover TL, Solnick JV. 2013. Functional plasticity in the type IV secretion system of Helicobacter pylori. PLoS Pathog. 9:e1003189.

68. Altman E, Chandan V, Li J, Vinogradov E. 2011. Lipopolysaccharide structures of Helicobacter pylori wild-type strain 26695 and 26695 HP0826::Kan mutant devoid of the O-chain polysaccharide component. Carbohydr Res. 346:2437–44.

69. Ma B, Audette GF, Lin S, Palcic MM, Hazes B, Taylor DE. 2006. Purification, kinetic characterization, and mapping of the minimal catalytic domain and the key polar groups of Helicobacter pylori alpha-(1,3/1,4)-fucosyltransferases. J Biol Chem. 281:6385–94.

70. Sun HY, Lin SW, Ko TP, Pan JF, Liu CL, Lin CN, Wang AH, Lin CH. 2007. Structure and mechanism of Helicobacter pylori fucosyltransferase. A basis for lipopolysaccharide variation and inhibitor design. J Biol Chem. 282:9973–82.

71. Lin SW, Yuan TM, Li JR, Lin CH. 2006. Carboxyl terminus of Helicobacter pylori alpha1,3-fucosyltransferase determines the structure and stability. Biochemistry. 45:8108–16.

72. Eren AM, Esen OC, Quince C, Vineis JH, Morrison HG, Sogin ML, Delmont TO. 2015. Anvi’o: an advanced analysis and visualization platform for ’omics data. PeerJ. 3:e1319.

73. Falk P, Roth KA, Boren T, Westblom TU, Gordon JI, Normark S. 1993. An in vitro adherence assay reveals that Helicobacter pylori exhibits cell lineage-specific tropism in the human gastric epithelium. Proc Natl Acad Sci U S A. 90:2035–9.

74. Yang H, Smith RD, Chandler CE, Johnson JK, Jackson SN, Woods AS, Scott AJ, Goodlett DR, Ernst RK. 2022. Lipid A Structural Determination from a Single Colony. Anal Chem. 94:7460–65.

75. Niedermeyer TH, Strohalm M. 2012. mMass as a software tool for the annotation of cyclic peptide tandem mass spectra. PLoS One. 7:e44913.

76. Davis MR, Jr., Goldberg JB. 2012. Purification and visualization of lipopolysaccharide from Gram-negative bacteria by hot aqueous-phenol extraction. J Vis Exp.

77. Taylor JA, Bratton BP, Sichel SR, Blair KM, Jacobs HM, DeMeester KE, Kuru E, Gray J, Biboy J, VanNieuwenhze MS, Vollmer W, Grimes CL, Shaevitz JW, Salama NR. 2020. Distinct cytoskeletal proteins define zones of enhanced cell wall synthesis in Helicobacter pylori. Elife. 9:

78. Wright MH, Adelskov J, Greene AC. 2017. Bacterial DNA Extraction Using Individual Enzymes and Phenol/Chloroform Separation. J Microbiol Biol Educ. 18:

